# Rapid evaluation of habitat connectivity change to safeguard multispecies persistence in human-transformed landscapes

**DOI:** 10.1101/2023.11.23.568419

**Authors:** Jacqueline Oehri, Sylvia L.R. Wood, Eluna Touratier, Brian Leung, Andrew Gonzalez

## Abstract

Protecting habitat connectivity in fragmented landscapes is essential for safeguarding biodiversity and nature’s contributions to people. Following the Post-2020 Kunming-Montreal Global Biodiversity Framework (KM-GBF) of the Convention on Biological Diversity (CBD) there is a clear science-policy need to assess habitat connectivity and track its change over time to inform conservation planning.

In response to this need we describe an analytical, multi-indicator and multispecies approach for the rapid assessment of habitat connectivity at fine spatial grain and at the extent of an entire ecoregion. Out of 69 connectivity indicators we found through a literature review, we identified a key-set of nine indicators that align with the Essential Biodiversity Variables framework and that are suitable to guide rapid action for connectivity and conservation targets in the KM-GBF. Using these selected indicators, we mapped and evaluated connectivity change from 2011 to 2021 across the ecoregion of the St-Lawrence Lowlands in Quebec (∼30,000 km^2^) for seven ecoprofile species representing regional forest habitat needs. For the majority of these ecoprofile species, trends over the last decade indicate a decline in effective connected area and metapopulation carrying capacity, mainly via a division of large contiguous habitat into smaller fragments, whereas total habitat area largely remained unchanged.

These results highlight that trends in habitat area and connectivity are not necessarily correlated and the urgent need to conserve and restore connectivity in the St-Lawrence Lowlands, in order to meet regional targets under the KM-GBF. Our general approach enables a comprehensive evaluation of connectivity for regional spatial planning for biodiversity. We develop an R-tool to support this analysis and that can be extended to other conservation planning efforts for connectivity.

## Introduction

Safeguarding ecological connectivity, the ‘unimpeded movement of species and the flow of natural processes that sustain life on Earth’ (CMS, 2019) is essential for effective biodiversity conservation and has been incorporated as a central element of the targets of the Kunming-Montreal Post-2020 Global Biodiversity Framework (KM-GBF) of the Convention on Biological Diversity (CBD, 2021). Ecological connectivity (hereafter ‘connectivity’) encompasses both the capacity of a landscape to maintain viable routes for potential species movement through physically linked habitat patches (structural connectivity, Calabrese & Fagan, 2004) and the realized ability of species to move through the landscape (functional connectivity, Salgueiro et al., 2021; Tischendorf & Fahrig, 2000). A key challenge is finding the measures and methods for assessing both aspects of change in connectivity over time to guide spatial conservation planning.

There exists a multitude of indicators for monitoring the structural and functional facets of connectivity (Keeley et al., 2021) but how they compare and what they imply for conservation management is often not easy to interpret (Hanson et al., 2022; Lalechère & Bergès, 2021; Wood et al., 2022). To date, common definitions, goals and standards to measure and evaluate connectivity change have not been well-established and consequently, criteria for critical connectivity thresholds are often not operationalized in spatial conservation planning (Beger et al., 2022; Ward et al., 2020; Wood et al., 2022). For example, the KM-GBF currently lacks an agreed headline indicator for monitoring connectivity and guiding action for 2030, although some component indicators have been proposed in the GBF’s monitoring framework (COP decision 15/5). Establishing a set of indicators and guidelines for their interpretation is an urgent need given that action targets are set for 2030 (CBD, 2021). Additionally, most connectivity indicators are difficult to interpret biologically in terms of the long-term outcomes for the persistence (e.g., extinction risk) for multiple species over a range of scales (Wood et al., 2022). The situation exists in part because rapid assessments of the connectivity needs of a wide range of taxa across scales remains both a theoretical and computational challenge (Wood et al., 2022).

A recent review by (Wood et al., 2022) highlights the potential of modelling multispecies connectivity either using carefully constructed “ecoprofiles” (Opdam et al., 2008), where a single species is selected or a generic species with composite trait values is created to represent the movement and habitat needs of a particular set of species or species group (Brodie et al., 2015) or, alternatively, “multiple focal species” approaches (Albert et al., 2017; Correa Ayram et al., 2018; Meurant et al., 2018), where landscape connectivity is modeled separately across a set of species with diverse ecological traits and their complementary connectivity priorities are identified and prioritized *post hoc*.

We conducted a literature review to identify commonly applied connectivity indicators that are suited for multispecies assessments, and that align with the criteria for Essential Biodiversity Variables (EBV) defined by the Group on Earth Observations Biodiversity Observation Network, such that they are feasible, (i.e. allow the connectivity monitoring for multiple species with minimal data inputs) are scalable, are sensitive to temporal change, and are relevant for biodiversity conservation targets (Jetz et al., 2019; Pereira et al., 2013). We searched for metrics that can be computed from relatively simple habitat distribution maps (Beger et al., 2022; Ward et al., 2020) to ensure wide scale applicability. The selected indicators include proxies of connectedness at pixel- and patch-level, as well as estimates of habitat-network connectivity at landscape-level. We also included metapopulation capacity as an indicator of potential long-term species persistence (Hanski & Ovaskainen, 2000; Hanski, 1994; Huang, Pimm, & Giri, 2020; Schnell et al., 2013; Stott et al., 2010; Strimas-Mackey & Brodie, 2018). To facilitate the interpretation and use of these indicators, we compare and contrast their characteristics and assess their correlation and sensitivity to changing habitat amount and fragmentation using a set of simulated landscapes (Saura & Martínez-Millán, 2000; Sciaini et al., 2018).

Using this set of selected multispecies connectivity indicators we developed a tool coded in R, “Rapid Evaluation for Connectivity Indicators and Planning” (RE-Connect, https://github.com/xxx/RE-Connect, Figure 1), and we used it to assess habitat connectivity and its temporal change from 2011 to 2021, for seven ecoprofile species representative of regional forest habitat and connectivity needs (Albert et al., 2017; Meurant et al., 2018; Rayfield et al., 2016) in the ecoregion of the St-Lawrence Lowlands in Quebec, Canada (Figure 2).

**Figure 1.**
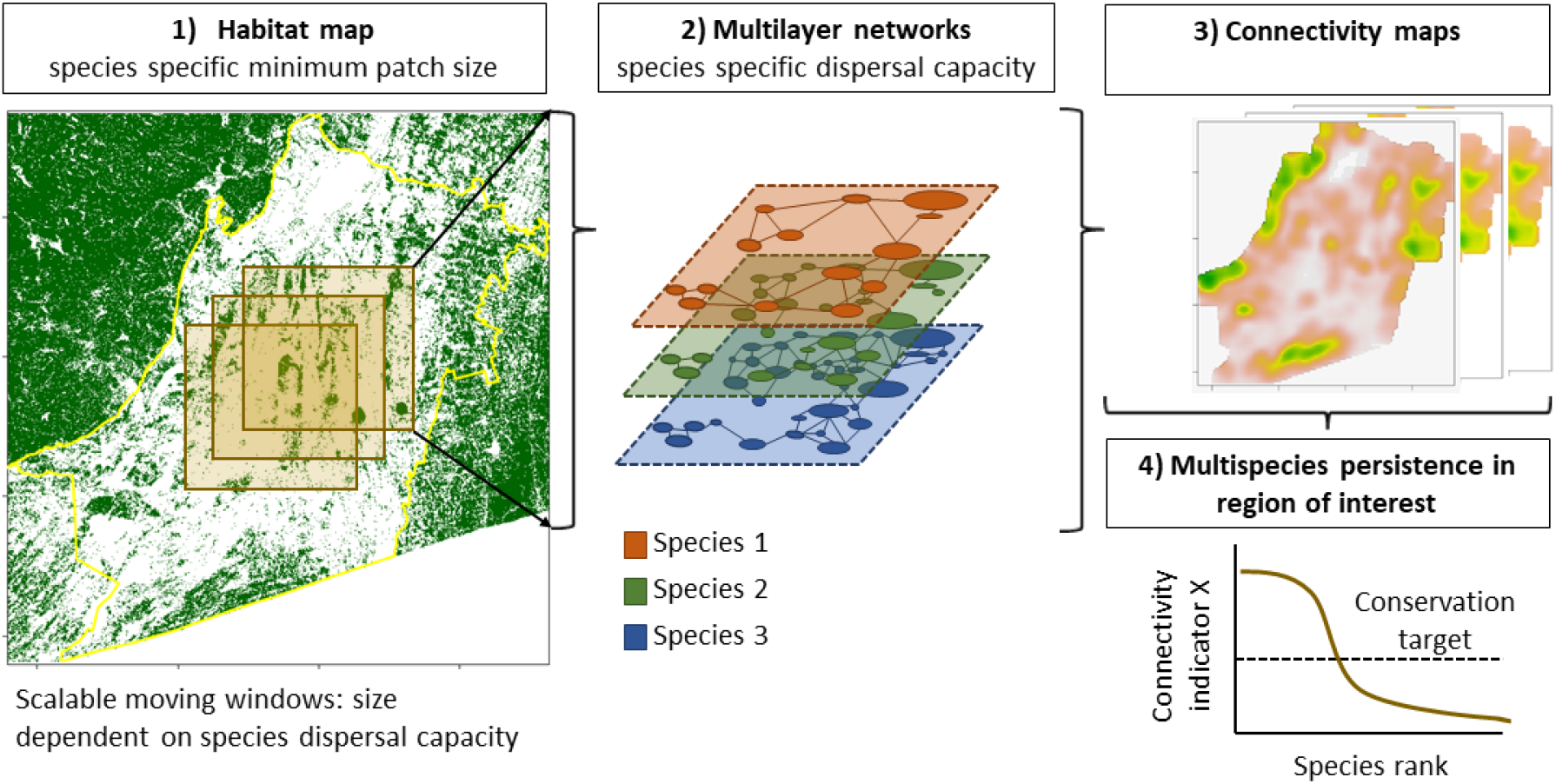
Steps for conducting a Rapid Evaluation of Multispecies Connectivity using the RE-Connect approach and R-tool. **1)** Input data for the RE-Connect tool is a species (or ecoprofile) specific, binary habitat map that can be determined *a priori* (e.g. from described habitat needs) or *a posteriori* (e.g., from species distribution models). **2)** Multiple habitat maps can be “stacked” and multilayer habitat networks can be extracted, in which links between habitat patches depend on species-specific dispersal capacities. Multilayer habitat networks can be computed in moving windows of relevant size and variable spatial overlap. **3)** Multiple connectivity indicators can be computed simultaneously for multiple species in the moving windows. Moving window results can be aggregated into mosaics, i.e. coherent maps of connectivity at pixel-level, patch-level or landscape-level for the species of interest. **4)** The resulting maps can be used to evaluate multiple connectivity indicators for the multiple species with regard to a defined target- or minimum threshold.

**Figure 2.**
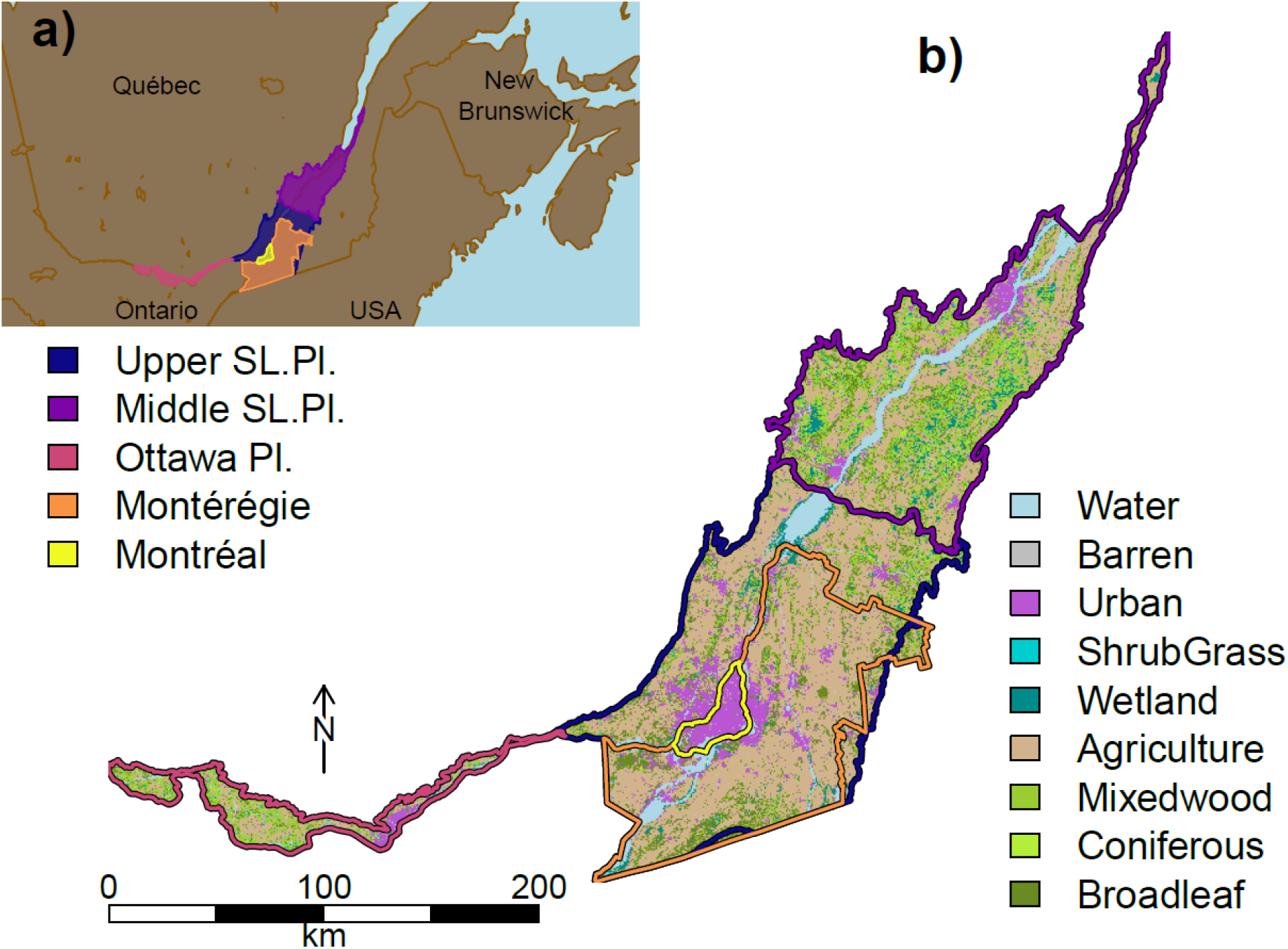
St-Lawrence Lowlands study region. **a)** The St-Lawrence Lowlands ecoregion is a priority landscape within the “Pan-Canadian Approach to Transforming Species at Risk Conservation in Canada” (ECCC, 2018; Jobin et al., 2020), spans ∼30,000 km^2^ and is located in Eastern Canada. **b)** The St-Lawrence Lowlands consists of three subregions: The Upper St-Lawrence Plain (including the regions of interest Montreal and Montérégie), the Middle St-Lawrence Plain and the Ottawa Plain and is composed predominantly of agricultural, urban and mixedwood land-cover classes.

The RE-Connect R-tool is based on recently developed R-packages (Csardi & Nepusz, 2006; van Etten, 2017; Godínez-Gómez, O. & Correa Ayram C.A., 2020; Hesselbarth et al., 2019) and R-functions in (Huang et al., 2020; Strimas-Mackey & Brodie, 2018). It allows for the assessment of multiple connectivity indicators simultaneously for multiple species (cf. multilayer habitat networks in Hartfelder et al., 2020), using simple habitat distribution maps and a parallel implementation of moving windows that are adjustable in form, size and spatial overlap (Drielsma & Ferrier, 2009; Hughes et al., 2023).

The St-Lawrence Lowlands study region has unique biogeographic characteristics, significant biodiversity and is affected by urbanization, agricultural expansion and habitat fragmentation, especially of forests and wetlands (Albert et al., 2017; Dupras et al., 2016; Lucet & Gonzalez, 2022; Mitchell et al., 2015; Rayfield et al., 2021). This ecoregion is one of eleven priority places within the “Pan-Canadian Approach to Transforming Species at Risk Conservation in Canada” (ECCC, 2018; Jobin et al., 2020).

Based on the results of our study, we identify areas with high priority for connectivity conservation. By assessing connectivity for multiple species with diverse habitat and movement needs, in a way that allows for a meaningful evaluation of long-term species persistence, we further support conservation management that safeguards biodiversity in the long-term.

## Methods

### Multispecies connectivity indicators literature review

We conducted a literature review according to the PRISMA approach to identify indicators suitable for the rapid assessment of multispecies connectivity across scales and regions of interest (Appendix S1, Liberati et al., 2009).

Specifically, on January 24th, 2022, we searched all journal articles published since January 1, 2000 on the Scopus and Web of Science databases using the following search string: habitat OR ecosystem OR “protected area” AND connectivity AND indicator OR metric OR indice OR index AND (multi OR several OR group) NEAR/3 species OR community AND conservation OR management OR monitoring OR planning AND biodiversity. This search led to a total of 1028 unique articles across queried databases. In a first screening step, we excluded all articles that did not mention either connectivity or monitoring in their title, leading to a total of 431 articles. In a second screening step, we excluded articles that 1) were not concerned with habitat connectivity, 2) were not primary research articles, 3) did not use any indicator of connectivity, or 4) were focused on a single species only, however articles that used a single species as umbrella species were kept. We reviewed the resulting 208 articles, identified the connectivity indicators they used, as well as the ecosystem types and taxa they were concerned with. Articles that did not align with our earlier criteria were removed, leaving a final set of 171 articles.

These 171 articles featured 69 indicators applied to quantify connectivity for multiple species. To distill a comprehensive subset of these indicators suitable for multispecies connectivity monitoring, we evaluated their correspondence with criteria established for the Essential Biodiversity Variable’s framework of GEO BON (Jetz et al., 2019; Pereira et al., 2013; geobon.org/ebvs/). Specifically, we aimed at synthesizing a set of indicators that i) are feasible, i.e. are commonly applied in the literature and can readily be computed from available data, ii) are relevant, i.e. allow for a coherent interpretation and alignment with biodiversity targets, iii) are scalable, i.e. computable across spatial scales and generalizable across a range of species and ecosystems, iv) are sensitive to connectivity change over time, and v) are complementary with regard to scale and interpretation (Appendix S2).

Based on our literature review, we summarise a set of nine connectivity indicators that are well suited for the multispecies contexts, are commonly applied in the scientific literature, offer a coherent interpretation, are relevant for conservation targets and can be efficiently computed based on simple habitat distribution maps across many species and environmental contexts (Table 1).

**Table 1.** Selected set of connectivity indicators^1^.

| N<br>r | Abbreviation | Description | Formula | Unit [value range] | Interpretation | Scale and distance type |
| --- | --- | --- | --- | --- | --- | --- |
| 1 | **MPC** | Metapopulation Capacity (Hanski & Ovaskainen, 2000; Hanski, 1994; Schnell et al., 2013) | $MPC = \lambda_m$ $m_{ij} = \begin{cases} f(d_{ij})a_ja_i^x & i \neq j \\ a_ja_i^x & i = j \end{cases}$<br>, where $\lambda_m$ is the leading eigenvalue of a square 'landscape matrix' $m$, in which elements $m_{ij}$ reflect rates of change for the occupancy of patches $i$ ($p_i$) as a function of patch attributes (often patch area in $m^2$, $a_i$ and $a_j$), a dispersal probability function of interpatch distance $f(d_{ij})$ and an extinction probability constant $x$ (commonly set to 0.5). | no unit<br>[0, $\lambda_{max}$]<br>, where $\lambda_{max}$ corresponds to the leading eigenvalue of the maximum landscape matrix, i.e. where the rates of change of patch occupancy is 1 for all patches. | -metapopulation carrying capacity, based on area and connectance of habitat<br>-focus on potential long-term species persistence (Hanski & Ovaskainen, 2000; Hanski, 1994; Schnell et al., 2013) | - landscape<br>- Euclidean and resistance |
| 2 | **ECA** | Equivalent Connected Area index (Saura et al., 2011), based on the probability of connectivity index (PC; Saura & Pascual-Hortal, 2007) | $ECA = \sqrt{\sum_{i=1}^n \sum_{j=1}^n a_i a_j p_{ij}^*}$<br>, where $n$ is the total number of habitat patches in the landscape, $a_i$ and $a_j$ are attributes of habitat patches $i$ and $j$ (commonly area) and $p_{ij}^*$ is the maximum product probability of dispersal between patches $i$ and $j$. | unit of $a$ (e.g. area) [0, $AI$]<br>, where $AI$ is the attribute (e.g. area) of the entire landscape. | -size of a single habitat patch that would provide the same probability of connectivity than the actual habitat pattern in the landscape (Saura et al., 2011)<br>-focus on connectivity of habitat, and potential species dispersal | - landscape<br>- Euclidean and resistance |
| 3 | **ECA<sub>Ap</sub>** | Fraction of habitat that is connected (ECA divided by the total habitat area) | $ECA_{Ap} = \frac{ECA}{Ap}$<br>, where $Ap$ is the total area of habitat in a given landscape. | no unit<br>[0, 1] | -size of equivalent connected area (ECA) relative to maximum possible with a given habitat area<br>-focus on underused connectivity potential | - landscape<br>- Euclidean and resistance |
| 4 | **ECA<sub>AI</sub>** | Fraction of habitat that is connected (ECA) divided by the total landscape area, cf. ProtConn index (Saura et al., 2017) | $ECA_{AI} = \frac{ECA}{AI}$ , where AI is the total area of a given landscape. | no unit<br>[0,1] | -size of equivalent connected area (ECA) relative to maximum possible in a landscape<br>-focus on underused connectivity potential | - landscape<br>- Euclidean and resistance |
| 5 | **BC** | Betweenness centrality (Albert et al., 2017; Brandes, 2001; Freeman, 1978) | $BC_v = \sum_{i \neq j, i \neq v, j \neq v} \frac{g_{ivj}}{g_{ij}}$ , where g<sub>ivj</sub> is the total number of shortest paths between each pair of habitat patches ij going through a focal patch v in the habitat network. g<sub>ij</sub>, is the total number of all the shortest paths between each pair of habitat patches ij in the habitat network. | no unit<br>$[0, \frac{(n-1)(n-2)}{2}]$<br>, where n corresponds to the number of patches in the landscape. | -number of shortest paths between each pair of habitat patches passing through a focal patch<br>-focus on the degree to which a habitat patch is a stepping stone at short and long distances in the habitat network (Keeley et al., 2021) | - patch<br>- Euclidean and resistance |
| 6 | **ND** | Node Degree (Minor & Urban, 2008) | $ND_v = \sum_{j \neq v} e_{vj}$ $e_{vj} = \begin{cases} 1 & p_{vj}^* > c \\ 0 & p_{vj}^* \leq c \end{cases}$ , where e<sub>vj</sub> denotes a habitat patch j connected to the focal patch v. A habitat patch can be considered connected (e<sub>vj</sub> = 1) if p<sub>ij</sub><sup>*</sup>, the maximum product probability of dispersal between patches j and v is greater than a threshold probability c. | no unit<br>[0,n-1]<br>, where n corresponds to the number of patches in the landscape. | -number of other habitat patches connected to a focal patch<br>-focus on the number of patches reachable within a given distance (Minor & Urban, 2008), the degree to which a patch is a stepping stone at short distances in the habitat network | - patch<br>- Euclidean and resistance |
| 7 | **I<sub>v</sub>** | Patch importance for landscape-level connectivity indicators (Saura & Pascual-Hortal, 2007) | $I_v = \frac{I - I'}{I} \times 100$ , where I is an index value when habitat patch v is present and I' is the same index value when patch v is not present in the landscape. | no unit<br>[0,100] | -the degree to which a patch contributes to the connectivity value of the habitat network | - patch<br>- Euclidean and resistance |
| 8 | **MPC<sub>i</sub>** | Patch importance for metapopulation capacity: leading eigenvector at position i (Hanski & Ovaskainen, 2000) | $MPC_i = \lambda_{mi}$ $m_{ij} = \begin{cases} f(d_{ij})a_j a_i^x & i \neq j \\ a_j a_i^x & i = j \end{cases}$<br>, where $\lambda_{mi}$ is the leading eigenvector at position i, for patch i of a square 'landscape matrix' m, in which elements $m_{ij}$ reflect rates of change for the occupancy of patches i ($p_i$). $a_i$ and $a_j$ are patch attributes (often patch area in $m^2$), $f(d_{ij})$ is a dispersal probability function of interpatch distance x denotes an extinction probability constant (commonly set to 0.5). | no unit<br>[0, $\lambda_{max}$]<br>, where $\lambda_{max}$ corresponds to the leading eigenvalue of the maximum landscape matrix, i.e. where the rates of change of patch occupancy is 1 for all patches. | -the degree to which a patch contributes to the metapopulation capacity value of the habitat network | - patch<br>- Euclidean and resistance |
| 9 | **invCR** | Omnidirectional inverse cumulative resistance (Albert et al., 2017) | $\frac{1}{CR} = \frac{n}{\sum_{k=1}^n CR_k}$<br>, where n is the number of pairs of locations selected in a given landscape and $CR_k$ is the cumulative resistance of the shortest paths between each pair of locations k. | variable unit<br>(0, $+\infty$) | -ease of traversability of the landscape<br>-focus on contribution to global, omnidirectional connectivity | - pixel and landscape<br>- resistance |

<sup>1</sup>This set of connectivity indicators has been identified through a literature review and selection criteria aligned with the Essential Biodiversity Variables (Jetz et al., 2019; Pereira et al., 2013) framework (Appendix S1 and S2). These indicators can be calculated on the basis of simple habitat or protected area distribution maps, species-specific habitat needs and dispersal capacities. Distance type: can be Euclidean or scaled by a resistance surface (Albert et al., 2017; Chubaty, Galpern, & Doctolero, 2020; Shahnasari et al., 2019). Scale: the scale at which indicator is typically calculated (pixel-, patch-, or landscape-level). Landscape-level indicators can also be calculated at patch level using the “patch importance for landscape-level connectivity indicators” computation ( $I_v$ ), or in the case of metapopulation capacity, the dominant eigenvector value for patch i (**MPC<sub>i</sub>**). Note that patch-level indicators can usually be aggregated at the landscape-level, using any aggregation function of interest (e.g. mean, range, variation). Note that our RE-Connect approach is not limited to these indicators but can be implemented with any indicator that is computable on the basis of habitat, protected area or resistance surface maps.

### Study region, species and habitat selection

We focused on the St-Lawrence Lowlands ecoregion of Quebec, Canada as defined by the Quebec Ecological Reference Framework (CERQ, Direction de l’expertise en biodiversité, 2018, Figure 2), consisting of the upper St-Lawrence Plain (including the areas of Montréal and the Montérégie, 17,315 km^2^), the middle St-Lawrence Plain (11,156 km^2^), the Ottawa Plain (2,223 km^2^), as well as the Montérégie (9,514 km^2^) and Montréal subregions of interest (627 km^2^). The St-Lawrence Lowlands harbors a relatively large fraction of Quebec’s biodiversity and is home to more than 55 species at risk (Tardif, Lavoie, & Lachance, 2005). At the same time, more than 4 million people (more than half of Quebec’s human population) live in this ecoregion, and anthropogenic activities that compromise the integrity of ecosystems are extensive, driven by urbanization, intensive agriculture and forestry, as well as the presence of invasive species (Jobin et al., 2020).

To model multispecies connectivity, we focused on seven species that depend on forest habitat and connectivity, and which were used by previous research (Albert et al., 2017; Meurant et al., 2018; Rayfield et al., 2016): American marten (*Martes americana*), black bear (*Ursus americanus*), northern short-tailed shrew (*Blarina brevicauda*), red-back salamander (*Plethodon cinereus*), wood frog (*Rana sylvatica*), ovenbird (*Seiurus aurocapilla*) and barred owl (*Strix varia*, Table 2). These species are a representative subset (Meurant et al., 2018) of an initial set of 14 surrogate species identified in (Albert et al., 2017) that cover the connectivity needs of forest-dependent terrestrial vertebrates across the St-Lawrence Lowlands. We selected the same subset of five species identified by Meurant et al., (2018) but also included the ovenbird and barred owl in order to represent birds with distinct forest needs. These species can simultaneously be interpreted as surrogate as well as ecoprofile species (Wood et al., 2022), which represent a regionally relevant range of forest habitat needs and dispersal capacities (Albert et al., 2017). Habitat needs were identified by Albert et al. (2017) and reflect the type of forest each species depends upon (coniferous, broadleaf and mixed) as well as the minimum habitat patch area that sustains a reproductive pair of a species population (Albert et al., 2017). Additionally, we define species-specific dispersal capacity as the capacity to move from one habitat patch to another, and therefore use the mean gap-crossing distance (see Table 2 in Albert et al., 2017).

**Table 2.**
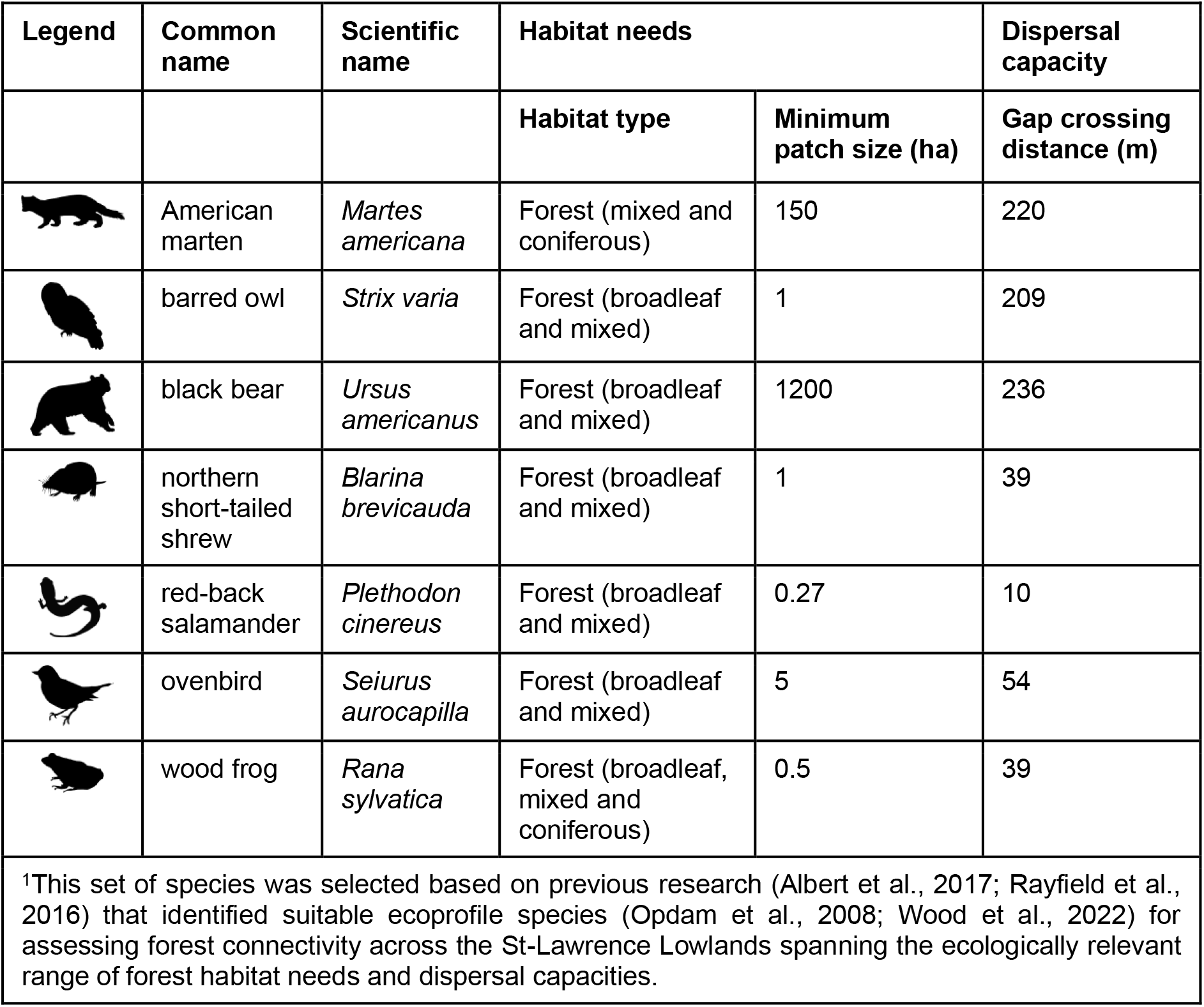
Selection of focal species for multispecies forest connectivity assessment^1^.

| Legend | Common name | Scientific name | Habitat needs |  | Dispersal capacity |
| --- | --- | --- | --- | --- | --- |
|  |  |  | Habitat type | Minimum patch size (ha) | Gap crossing distance (m) |
|  | American marten | *Martes americana* | Forest (mixed and coniferous) | 150 | 220 |
|  | barred owl | *Strix varia* | Forest (broadleaf and mixed) | 1 | 209 |
|  | black bear | *Ursus americanus* | Forest (broadleaf and mixed) | 1200 | 236 |
|  | northern short-tailed shrew | *Blarina brevicauda* | Forest (broadleaf and mixed) | 1 | 39 |
|  | red-back salamander | *Plethodon cinereus* | Forest (broadleaf and mixed) | 0.27 | 10 |
|  | ovenbird | *Seiurus aurocapilla* | Forest (broadleaf and mixed) | 5 | 54 |
|  | wood frog | *Rana sylvatica* | Forest (broadleaf, mixed and coniferous) | 0.5 | 39 |
| <sup>1</sup>This set of species was selected based on previous research (Albert et al., 2017; Rayfield et al., 2016) that identified suitable ecoprofile species (Opdam et al., 2008; Wood et al., 2022) for assessing forest connectivity across the St-Lawrence Lowlands spanning the ecologically relevant range of forest habitat needs and dispersal capacities. |  |  |  |  |  |

### Decadal change of multispecies connectivity across the St-Lawrence Lowlands

We assessed the state of habitat connectivity as captured by all our indicators at landscape-level and patch-level (i.e., except the invCR indicator in Table 1, to reduce complexity of our analysis), for the seven species (Table 2) across the St-Lawrence Lowlands ecoregion for the years 2011 and 2021, using the RE-Connect tool (Figure 1).

To do so, we derived species-specific, binary habitat maps (i.e. gridded spatial data with location of habitat vs. non-habitat areas) from land-cover information provided by the Annual Crop Inventory data layer (30 m resolution, Agriculture and Agri-Food Canada, 2023) for the years of 2011 and 2021. In particular, we extracted the land-cover classes "Coniferous", "Broadleaf", "Mixedwood" and combined these with minimum habitat patch size criteria for each species to classify discrete habitat patches. In these species-specific habitat networks, we determined the species-specific dispersal probabilities among habitat patches by a negative exponential kernel (Hartfelder et al., 2020; Moilanen & Nieminen, 2002; Strimas-Mackey & Brodie, 2018):

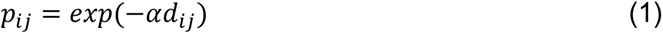

, where **α** is the inverse mean gap crossing distance (Table 2, cf. Hanski & Ovaskainen, 2000; Hartfelder et al., 2020; Moilanen & Nieminen, 2002) and *dij* is the Euclidean edge-to-edge distance between patch *i* and patch *j* in the habitat network.

Using our RE-Connect tool, we computed connectivity indicators in square moving windows with a side length of 8,700 m and an area of ∼75 km^2^, similar to previous research (Huang et al. 2020, Strimas-Mackey & Brodie, 2018). To build seamless landscape results across our moving window analysis we chose a window overlap of 1,500 m. Using these settings, we generated species-specific maps of landscape-level connectivity indicators, by averaging the connectivity values in ∼75 km^2^ overlapping moving window landscapes for each pixel at a 1.5 km effective resolution. Similarly, we generated species-specific maps of patch-level connectivity indicators, by computing the weighted mean values for each patch covered by overlapping moving window landscapes (weighted by the respective area patches covered in the overlapping moving-window landscapes).

To quantify the decadal change in the spatial distribution of species-specific connectivity, we subtracted RE-Connect results for 2011 from those calculated for 2021, for each species and connectivity indicator separately. Next, we averaged these species-specific maps of connectivity change to generate a map of the spatial distribution of ensemble gains and losses in connectivity across the St-Lawrence Lowlands for each of the selected connectivity indicators separately (Figure 3).

**Figure 3.**
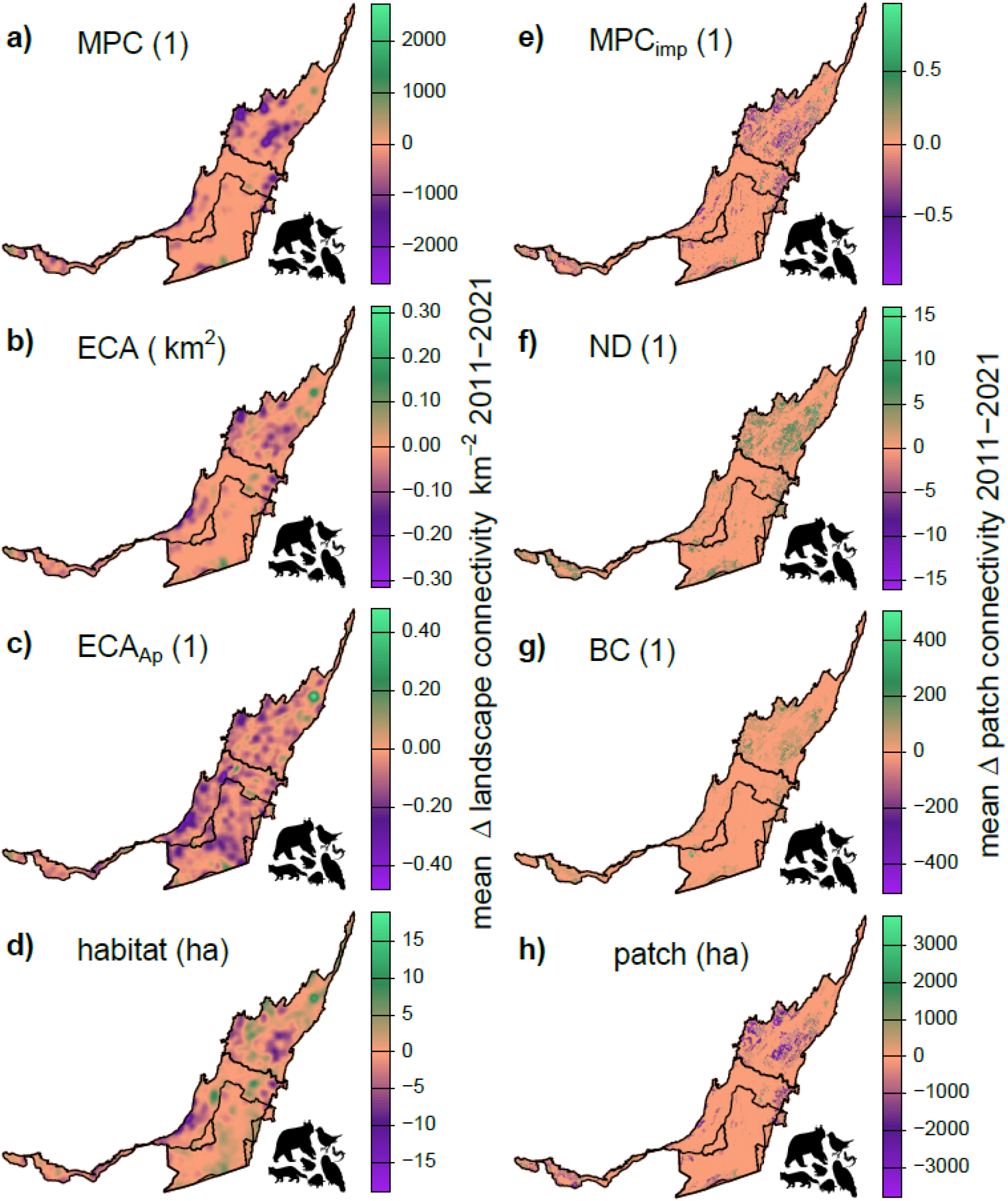
Decadal change (2011-2021) in multispecies connectivity indicators at the landscape-level (a-d) and patch-level (e-h) across the St-Lawrence Lowlands as calculated by RE-Connect. **a)** MPC: metapopulation capacity, **b)** ECA: equivalent connected area index (km^2^), **c)** ECA_Ap_: fraction of habitat that is connected, **d)** habitat area in hectares, **e)** MPC_imp_: metapopulation capacity patch importance, **f)** ND: node degree of focal patches, g) BC: betweenness centrality of focal patches, **h)** area of focal patches in hectares. We scaled connectivity values by the area of the RE-Connect moving window size (8,700^2^ = 75.69 km^2^) if necessary, i.e. in the cases of MPC, ECA and habitat area. See Table 1 for more detailed explanations of connectivity indicators.

In a supplementary analysis, we provide ensemble connectivity maps for the year 2021 (Appendix S4), which resulted from averaging species-specific connectivity maps in 2021 that were previously normalized using the feature scaling approach:

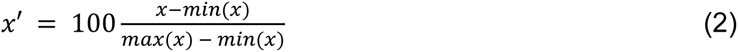

, where *x* is the vector of all values contained in the respective connectivity maps.

This normalization was done to focus on the relative magnitude of connectivity values and to give each species equal weight when aggregating *post hoc*.

### Evaluating multispecies connectivity change for species persistence

In order to shift the focus from a spatial assessment to an actionable evaluation of connectivity across species, we used a species-rank approach (Chowdhury et al., 2023; Hartfelder et al., 2020; Silvestro et al., 2022) to rank connectivity indicator change values across five CERQ subregions of interest in the St-Lawrence Lowlands (Direction de l’expertise en biodiversité, 2018; Figure 4).

**Figure 4.**
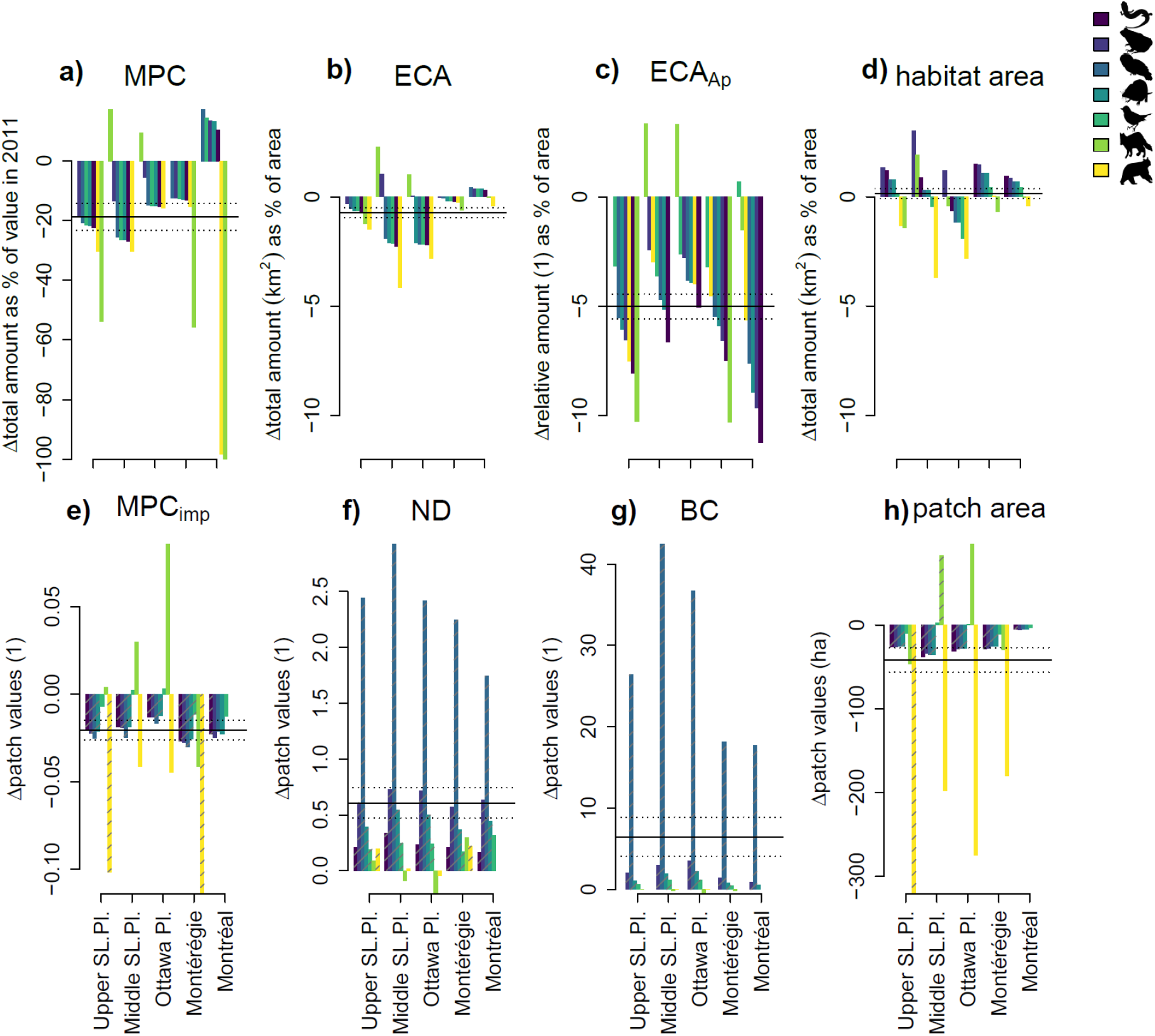
Decadal change (2011-2021) of multispecies connectivity indicators at landscape-level (a-d) and patch-level (e-h) for 5 different regions in the St-Lawrence Lowlands by species. **a)** MPC: metapopulation capacity, **b)** ECA: equivalent connected area index, **c)** ECAAp: fraction of habitat that is connected, **d)** habitat area: species-specific area of habitat, **e)** MPCimp: metapopulation capacity patch importance, **f)** ND: node degree of focal patches, **g)** BC: betweenness centrality of focal patches, **h)** patch area in hectares. Horizontal bars indicate the mean and s.e. of change across all regions and species. Hashed bars in e-h) indicate significant changes (p<0.05) in patch-connectivity magnitudes as indicated by two-sided Welch’s t-tests comparing patch values in 2021 and 2011. See Table 1 for more detailed explanations of connectivity indicators. See Appendix S6 and S7 for more details.

In particular, in the cases of landscape-level metapopulation capacity (MPC), equivalent connected area (ECA) and habitat area, we scaled the species-specific, RE-Connect derived connectivity change values by dividing by the area of the moving window size (8,700m × 8,700m = 75.69 km^2^). Then, we multiplied these landscape-level connectivity change values by the area they cover, and consequently summed these values to assess the total amount of connectivity change for each subregion of interest.

In the case of the patch-level connectivity indicators of metapopulation capacity patch importance (MPC_imp_), node degree (ND) and betweenness centrality (BC), as well as patch size (patch area), we extracted the distribution of values for each species and subregion of interest for the years 2011 and 2021. We then assessed the decadal change of patch-connectivity magnitudes by comparing connectivity magnitudes of all habitat patches in 2021 to those in 2011 using two-sided Welch’s unequal variances t-tests. We used Welch’s t-tests because they are more robust than Student’s t-tests when the two samples have unequal variances and/or unequal sample sizes, as was the case for our habitat patch values in 2021 and 2011, respectively (Ruxton, 2006).

In a supplementary analysis, we summarized and ranked connectivity results for each ecoprofile species within five subregions of interest for the year 2021 (Appendix S5).

### Comparing multispecies connectivity in simulated and real-world landscapes

To facilitate interpretation and use of the selected key-set of connectivity indicators, we generated a reference set of simulated landscapes orthogonal in their gradients of habitat amount and fragmentation using the “random-cluster” algorithm (Saura & Martínez-Millán, 2000) and implemented in the NLMR R-package (Sciaini et al., 2018). Specifically, we generated five landscape replicates (250 x 250 cells) along a gradient of habitat amount (0.1, 0.2, 0.3, 0.4, 0.5, 0.6, 0.7, 0.8 and 0.9 fractional cover in the landscape, parameter *Ai* in Saura & Martínez-Millán (2000), and habitat fragmentation (clumping factor 0.1, 0.2, 0.3, 0.4, 0.5 and 0.6, parameter *p* in Saura & Martínez-Millán (2000). In the resulting set of 5 x 9 x 6 = 270 simulated reference landscapes, we computed patch- and landscape-level connectivity indicators using our RE-Connect tool for five dispersal capacities – **α** for a negative exponential distribution [Eq. 1 above]. **α** values ranged from the inverse of 1 cell unit to 355 cell units, whereby 355 is slightly larger than the diagonal of the simulated landscapes (250 × 250 cells). Compared with landscape size, a gap crossing distance of 6.8 cells is comparable with the maximum gap crossing distance among our selected species (236 m, Table 2) in the moving window landscapes of ∼75km^2^ (i.e. both gap crossing distances correspond to ∼1/36 of the landscape side length). Using these 270 reference landscapes, we assessed the Pearson correlations among connectivity indicators (Figure 5a) for a gap crossing distance of 6.8 units.

**Figure 5.**
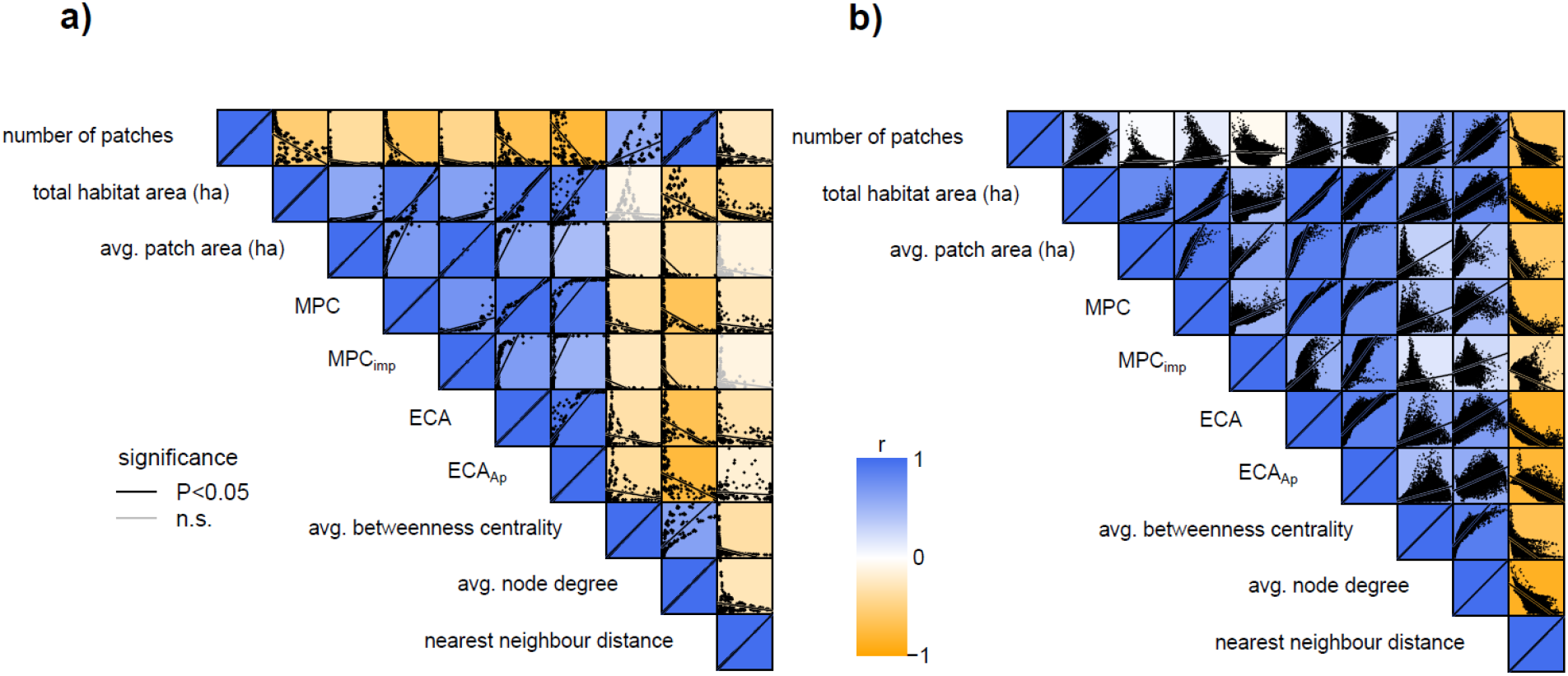
Pearson correlations among multiple connectivity indicators in simulated landscapes (a) and real-world landscapes in the St-Lawrence Lowlands (b). Number of patches: number of habitat patches in the landscape, total habitat area: amount of habitat area in landscape in hectares, MPC: metapopulation capacity, MPC_imp_: metapopulation capacity patch importance, ECA: Equivalent Connected Area index, ECA_Ap_: Fraction of habitat that is connected, avg. betweenness centrality: average number of shortest paths between all pairs of habitat patches that go through focal patches, avg. node degree: average number of habitat patches connected to focal patches, nearest neighbor distance: average Euclidean distance to nearest neighbor for focal patches in the landscape. See Table 1 for more detailed explanations of connectivity indicators.

To assess how the results obtained with the simulated landscapes relate to the real-world landscapes across the St-Lawrence Lowlands, we repeated the correlation analysis using a random sample of 10,000 cells from the ensemble connectivity maps we generated for the year 2021 using our RE-Connect tool (see Methods above, Figure 5b, Appendix S4). Finally, we assessed the sensitivity of connectivity indicators to changes in habitat area (Appendix S3a-d), and habitat fragmentation as approximated by the number of habitat patches (Appendix S3e-h) across all gap crossing distances from 1 to 355 cell units. Specifically, we estimated and predicted relationships among connectivity indicators and habitat area (and fragmentation) using a smooth spline function (npreg R-package, Helwig, 2021). This analysis allowed us to capture non-linearities in relationships among connectivity indicators and habitat area (or fragmentation), as well as assessing the effect of different gap crossing dispersal capacities (Appendix S3).

## Results

### Decadal change of multispecies connectivity across the St-Lawrence Lowlands

We assessed the spatial distribution of decadal patch- and landscape-level connectivity change for each species and indicator separately, by calculating the difference between 2021 and 2011. We then averaged results over all species to create a map of multispecies connectivity change for each indicator (Figure 3).

At the landscape-level, decreases in average multispecies metapopulation capacity (MPC) and equivalent connected area (ECA) are more abundant than gains across the St-Lawrence Lowlands (MPC: losses in 21,342 km^2^, gains in 9,663 km^2^, range: -2,727-1,101 km^-2^; ECA: losses in 18,876 km^2^, gains in 12,130 km^2^, range: -0.31-0.20 km^2^·km^-2^), especially in the Middle St-Lawrence Plain (Figure 3a,b). Similarly, decreases in the fraction of effectively connected habitat (ECA_Ap_, Figure 3c) prevail and can reach up to 48% (losses in 24,675 km^2^, gains in 6,331 km^2^, range: -0.44-0.48 km^-2^), whereas habitat area changes are more variable (losses in 12,510 km^2^, gains in 18,495 km^2^, range: -19-14 ha·km^-2^) across the St-Lawrence Lowlands ecoregion (Figure 3d).

Despite the declines described above, in 2021, the Middle St-Lawrence and Ottawa Plain subregions harbor the highest values of MPC, ECA, ECA_Ap_ and habitat area, and low connectivity indicator values are predominantly found in the Upper St-Lawrence Plain subregion (Appendix S4).

Patch-level multispecies connectivity in 2021 (Appendix S4) and connectivity changes from 2011 to 2021 (Figure 3e-h) show a similar spatial distribution as those at the landscape-level. In particular, patch importance for metapopulation capacity (MPC_imp_) tended to decrease, especially in the Middle St-Lawrence Plain (losses in 6,091 km^2^, gains in 4,139 km^2^, range: - 0.96-0.95 per patch; Figure 3e). In contrast, betweenness centrality (BC) and node degree (ND) show large increases, in the case of BC up to 505 shortest paths between each pair of habitat patches passing through a focal patch (BC: losses in 1,650 km^2^, gains in 8,160 km^2^, range: -122-505 per patch; ND: losses in 2,048 km^2^, gains in 8,151 km^2^, range: -7-16 per patch; Figure 3f,g). Also, habitat patch size tended to decrease across the entire ecoregion of St-Lawrence Lowlands, sometimes up to 3,806 hectares per patch (losses in 6,273 km^2^, gains in 4,004 km^2^, range: -3,767-3,806 ha per patch; Figure 3h).

### Evaluating multispecies connectivity change for species persistence

We summed and ranked connectivity change for each indicator and ecoprofile species within five subregions of interest in the St-Lawrence Lowlands ecoregion (Figure 4).

Analyses of decadal changes in landscape-level connectivity indicators (MPC, ECA, ECA_Ap_), reveal declines for the majority of species and subregions of interest. For example, metapopulation capacity declined by 18.8 ± 4.5 % (mean and s.e.) of its average value in 2011 (4,991,781 units) across species and subregions (black horizontal line, Figure 4a). Relative metapopulation capacity declines were most pronounced for the Upper St-Lawrence Plain (on average by -27.0 % of its 2011 value) and for the black bear and American marten (on average by -38.0 % and -36.6 % of their respective 2011 values). However, relative metapopulation capacity increased in the case of the American marten in the Middle St-Lawrence and Ottawa Plains by 17.3 % and 9.3 % of its 2011 value, respectively. Metapopulation capacity increased in the Montreal area for all species except the black bear and the American marten.

A similar pattern is shown for the equivalent connected area index (ECA, Figure 4b), which declined on average by 0.7 ± 0.2 % (mean and s.e.) of the total area across subregions and species. Relative declines were most pronounced in the Middle St-Lawrence and Ottawa Plains (average loss of 1.3 % and 1.5 % of area) and for the black bear (average loss of 1.8 % of area). The fraction of connected habitat (ECA_Ap_, Figure 4c) showed declines across subregions and species, with an average decline equivalent to a complete loss of habitat connectivity in 5 ± 0.6 % (mean and s.e.) of the total area. Similar to the case of metapopulation capacity and that of equivalent connected area index, ECA_Ap_ increased in the case of American marten in the Middle St-Lawrence and Ottawa Plains, as well as for the ovenbird in Montreal. Contrasting the connectivity indicator changes, changes in habitat area (Figure 4d) varied across subregions and species and increased on average by 0.1 ± 0.2 % (mean and s.e.). Hence, patterns of habitat area and connectivity change do not overlap and are not correlated across the species of interest.

In the case of the current status (2021) of landscape-level multispecies connectivity (Appendix S5), metapopulation capacity (MPC) values are highest in the Middle St-Lawrence Plain, and are generally high for ecoprofile species with small patch size requirements, such as the wood frog, or species with larger gap crossing capabilities, such as the barred owl. In the case of the effectively connected area index (ECA), on average, 14.6% of the St-Lawrence Lowlands total area consists of species habitat that is effectively connected. An ECA of 30% is only achieved in the Ottawa Plain (Appendix S5b). Also, the percentage of habitat area is on average higher across subregions and species (average: 20.8%), as compared to the percentage of effectively connected habitat area (ECA, average: 14.6%).

Using two-sided Welch’s t-tests, we compared patch-connectivity values in 2021 and 2011 and found significant declines in average metapopulation capacity patch importance (MPC_imp_), and average habitat patch sizes, as well as significant increases in average node degree and betweenness centrality of habitat patches across most subregions and species of interest (Figure 4e-h).

Patch-level connectivity distributions in 2021 tend to be skewed, with predominantly low median values across all subregions and all species except the American marten and the black bear, which had higher median values due to larger minimum patch size requirements (Table 2, Appendix S5e-h).

### Comparing multispecies connectivity in simulated and real-world landscapes

We summarise a set of nine multispecies connectivity indicators, their characteristics and interpretation in Table 1. An analysis of indicator correlations across our 270 simulated landscapes revealed that metapopulation capacity (MPC) -based and equivalent connected area (ECA) -based indicators form a cluster that is positively correlated with habitat area and negatively correlated with fragmentation (i.e., the number of patches, Figure 5a). Hence, MPC and ECA show a very similar relationship with habitat area and fragmentation, although MPC is focused on long-term dispersal outcomes and is therefore less sensitive to differences in dispersal capacity compared to ECA (Appendix S3).

In contrast to MPC and ECA, average betweenness centrality and node degree tend to be negatively correlated with habitat area and positively correlated with the number of patches in the simulated landscapes. Interestingly, this correlation pattern was less pronounced in the 10,000 randomly sampled cells across the St-Lawrence Lowlands (RE-Connect ∼75km^2^-moving window results for the year 2021, Appendix S4), where all connectivity indicators tend to be positively correlated with habitat area and number of patches (Figure 5b). Further analyses of the simulated landscapes revealed important non-linearities in the relationships between habitat area (Appendix S3a-d), number of patches (Appendix S3e-h) and connectivity indices. For example, average landscape-level node degree shows a hump-shaped relationship with habitat area (Appendix S3d), with a peak around a habitat area of ∼ 1.5 ha, i.e. when habitat covers ∼24% of the total landscape area. Since in the simulated landscapes, the amount of habitat area varied between 10% and 90% of the total landscape area (cf. Methods), the negative relationship between habitat area and node degree dominated in the correlation analysis (Figure 5a). However, in the moving window landscapes of the St-Lawrence Lowlands ecoregion, median habitat cover ranged from 3% (American marten) to 27% (wood frog), depending on the species, and therefore, the positive relationship between habitat area and node degree prevailed in the correlation analysis (Figure 5b).

## Discussion

In this study, we summarised the characteristics of nine key-indicators of ecological connectivity and developed a scalable, generalizable analytical approach and R-tool (RE-Connect) to rapidly evaluate connectivity indicators for multiple species with different habitat needs in any region of interest. With this tool we analysed the change in forest connectivity for the period 2011-2021 in the St Lawrence Lowlands ecoregion. We found a decline in forest connectivity for multiple species, as measured by metapopulation capacity and effective connected area. This change arose because of the fragmentation of large contiguous habitat into smaller patches, while the total habitat area remained largely unchanged. Additionally, we found that for the year 2021, the St-Lawrence Lowlands harbor on average 14.6% effectively connected forest habitat area, which is about half the target value of 30% (cf. Target 3 in the Post-2020 Global Biodiversity Framework of the Convention on Biological Diversity; CBD, 2021). These findings offer guidance to Quebec’s planning for connectivity in the south of the province where human impacts are greatest.

### Multispecies connectivity indicators supporting the Global Biodiversity Framework

Our connectivity indicators support connectivity assessments in regions with moderate to high human disturbance (Keeley et al., 2021), where biodiversity is under intense land-use pressure. They were selected to align with criteria for Essential Biodiversity Variables (EBV), such that they are feasible, (i.e. allow the monitoring of connectivity for multiple species with minimal data inputs) are scalable, are sensitive to temporal change, and are relevant for biodiversity conservation targets (Jetz et al., 2019; Pereira et al., 2013). While a wide range of EBVs have already been established, an EBV-based connectivity indicator has yet to be incorporated into the EBV framework (geobon.org/ebvs/). We believe that the indicators selected here have the potential to support the implementation and continued monitoring of connectivity targets for multiple species at local, national and global scales, and as complementary indicators for the KM-GBF.

For example, at the landscape-level, metapopulation capacity (MPC) captures the potential long-term persistence of species across a landscape (Drielsma & Ferrier, 2009; Hanski & Ovaskainen, 2000; Hanski, 1994; Hanski et al., 2017; Schnell et al., 2013). Meanwhile, the effectively connected area index (ECA; Saura et al., 2011; Saura & Pascual-Hortal, 2007) is especially relevant for area-based connectivity targets (CBD, 2021; Ward et al., 2020). The fraction of the effectively connected area index (ECA_Ap_) can be used to identify areas with underused connectivity potential (i.e. where habitat exists but is not connected). Further indices at the patch-level can highlight the degree to which a habitat patch is a stepping stone for movement over short distances (i.e. node degree, ND) or short and long distances (betweenness centrality, BC) in the habitat network. Hence, the list in Table 1 is not exclusive, but covers connectivity dimensions at different levels of spatial resolution and extent (Fletcher et al., 2023).

Our analyses highlight the existence of clusters of correlated indicators, such as MPC and ECA, that are positively correlated with habitat area, and negatively correlated with betweenness centrality, node degree and number of patches in simulated landscapes (Figure 5a). However, as our connectivity change assessment across the St-Lawrence Lowlands showed, despite being in a correlated cluster, the trends in MPC and ECA did not reflect the trend of habitat area from 2011 to 2021 (Figure 4). Additionally, because of non-linear change in configuration arising from change in habitat area (or fragmentation) we expect measures of connectivity (e.g., node degree Appendix S3d), and their correlations, to change in real-world landscapes even with quite modest changes in total amount of available habitat. Hence, a combination of multiple connectivity indicators at different levels (e.g., patch and landscape), selected according to specific conservation needs might be best to robustly assess and evaluate observed connectivity patterns. We found the use of simulated reference landscapes provided a benchmark against which to identify and target thresholds for connectivity.

### Diverging trends in habitat area and connectivity in the St-Lawrence Lowlands

The St-Lawrence Lowlands ecoregion is characterized by decades of profound landscape changes via intensification of agriculture and forestry, as well as urbanization. Currently, around 40% of its territory is characterized by agriculture of annual crops, forests cover 24%, urban lands 12% and wetlands around 10%. A recent analysis of land-cover change in the last 25 years (Drapeau et al., 2019; Jobin et al., 2020; Regos et al., 2018) shows that the forest cover slightly increased, which matches the pattern we observed for the last decade in our study. However, our results show that habitat area and connectivity trends do not necessarily correlate: despite the slight average increase in total forest habitat area, losses in connectivity dominate for most of our assessed species. Losses in MPC and ECA were most prevalent for the black bear, which was the species with the largest minimum patch size requirements in deciduous and mixed forest habitats. Our findings of a trend in decreasing average patch size and increasing betweenness centrality uncovers the typical phenomenology of habitat fragmentation: the division of habitat area into smaller and more isolated fragments (Haddad et al., 2015).

These patterns suggest that area-based habitat conservation alone might not be enough to safeguard biodiversity in the St-Lawrence Lowlands (Maxwell et al., 2020). Subregions with strong recent connectivity declines, such as the Middle St-Lawrence Plain, as well as subregions where connectivity values are generally low, such as the Upper St-Lawrence Plain should be the focus of connectivity conservation action in the future. These subregions were also ranked as highly important for maintaining species connectivity between large wilderness areas (provincial parks) and represent critical migration corridors for species moving north with climate change (Rayfield et al., 2021). Hence, reconnecting fragmented landscapes in these subregions could contribute to the long-term persistence of threatened species and contribute to the Pan-Canadian Approach to Transforming Species at Risk Conservation in Canada (ECCC, 2018).

Our results are focused on the connectivity of forest habitats, although other important habitats of concern for biodiversity conservation in the ecoregion are undergoing declines in area and connectivity, such as wetlands, grasslands, as well as freshwater ecosystems (Jobin et al., 2020). Future connectivity analyses should include inter-ecosystem connectivity. Our RE-Connect approach and R-tool would work in this case as well.

### Towards an actionable evaluation of multispecies connectivity

Ranking species according to the values within and across connectivity indicators is a simple, coherent and generalizable way to quantitatively evaluate the state and temporal trend of multispecies connectivity while also retaining species-specific information across any region of interest (Chowdhury et al., 2023; Hartfelder et al., 2020; Silvestro et al., 2022).

To meaningfully evaluate species-specific connectivity values, an explicit formulation of conservation targets is necessary (Drielsma & Ferrier, 2009). Area-based conservation targets such as “30% by 2030” in the KM-GBF (CBD, 2021; Gurney et al., 2023) should also account for connectivity. To this end the effective connected area index (ECA) can be used to assess the number of species and regions that meet a minimum habitat area threshold of 30% whilst also being effectively connected (Appendix S5b).

Importantly, if area-based conservation efforts are to address the global biodiversity crisis, they must not only consider connectivity, but also species persistence (Maxwell et al., 2020). Identifying explicit value ranges of indicators that represent a “safe operating space” (Gonzalez et al., 2017; Steffen et al., 2015) for species persistence has been a notorious challenge (Bulman et al., 2007; Flather et al., 2011; Hanski et al., 2017). The original model of the metapopulation capacity (MPC) indicator implies that a metapopulation persists in a habitat network if MPC is greater than the ratio of colonization and extinction rate parameters (cf. extinction threshold ***δ*=*e*/*c*** in Drielsma & Ferrier, 2009; Hanski & Ovaskainen, 2000; Hanski et al., 2017; Schnell et al., 2013). Because colonization and extinction rate parameters are hard to estimate in nature, minimum MPC and ECA values for species persistence could be based on a minimum viable number of reproductive pairs in a population (Albert et al., 2017) or the habitat requirements for a minimum viable metapopulation size (Bulman et al., 2007; Drielsma & Ferrier, 2009; Taylor et al., 2016). Nevertheless, more research combining models with empirical data is needed to assess the uncertainty around adequate thresholds for long term species (or community) persistence or extinction risk (Bulman et al., 2007; Flather et al., 2011; Hanski et al., 2017). The metapopulation capacity indicator, calibrated with empirical data could offer a sound basis towards that goal and can be expanded to capture different aspects of population, species and community dynamics, such as the extrapolated metapopulation persistence time (Schnell et al., 2013) and the potential food-chain length (Wang et al., 2021).

### Overcoming common challenges of multispecies connectivity modelling and future directions with the RE-Connect tool

We used *a priori* definitions of species dispersal capacity and a negative exponential kernel based on Euclidean distances to approximate dispersal probability among habitat patches. Hence, our approach lies between quantifying the capacity of a landscape to foster potential species movement (structural connectivity; Calabrese & Fagan, 2004) and expected realized species movement (functional connectivity; Salgueiro et al., 2021; Tischendorf & Fahrig, 2000). Realized species movement does not only depend on the structure of habitat networks, but also on other external factors such as population dynamics (Chu & Claramunt, 2023), and the behavior of individuals (Nathan et al., 2008; Rayfield et al., 2023). To generate more realistic connectivity estimates, movement traits could be estimated from species traits (e.g. body mass or wing length; Chu & Claramunt, 2023; Hartfelder et al., 2020) and included in more sophisticated dispersal probability kernels with resistance-weighted distance (McRae et al., 2016; Rayfield et al., 2023) or even in trait-based movement models (Hirt et al., 2018). Alternatively, dispersal could be derived empirically from *realized* movement trajectories (e.g. via GPS tracking or camera traps; Tucker et al., 2018), but such data can be difficult and costly to obtain (Wood et al., 2022).

Data scarcity is also among the reasons why connectivity assessments are rarely empirically validated (Daniel et al., 2023; Foltête et al., 2012; Foltête et al., 2020; Lalechère & Bergès, 2021; Laliberté & St-Laurent, 2020; Wood et al., 2022). Combining multispecies connectivity assessments with species distribution models based on openly accessible data constitutes a promising avenue to empirically test the importance of connectivity for species movement and persistence (Curd et al., 2022; Daniel et al., 2023; Lalechère & Bergès, 2021; Van Moorter et al., 2023; Vasudev et al., 2015). The ensemble connectivity maps generated with our RE-Connect approach could be combined with such species distribution models.

Modeling connectivity across large spatial extents at fine resolution can rapidly become computationally demanding (Albert et al., 2017; Koen, Ellington, & Bowman, 2019; Santini, Saura, & Rondinini, 2016). One set of strategies to address this challenge is based on decreasing the spatial extent by splitting a large study area into smaller, distinct windows with overlap (Drielsma & Ferrier, 2009; Hughes et al., 2023; Koen et al., 2019; Landau et al., 2021) or without overlap (Strimas-Mackey & Brodie, 2018). Another set of strategies is related to decreasing the spatial resolution of the input data (Koen et al., 2019), and thereby effectively aggregating (and reducing the number) habitat patches (Albert et al., 2017). A third set of strategies is focused on removing habitat patches with attribute values transgressing a threshold (e.g. minimum habitat patch size; Albert et al., 2017). Our RE-Connect analytical tool can be adjusted to include all of these strategies. Recent advances highlight the possibility of a generalizable subsampling of habitat networks in order to estimate whole network properties (Song et al., 2022), with the limitation that estimates can only be assessed at the whole network scale.

In the current study, we assessed connectivity at a single scale (moving windows of ∼75 km^2^). However, connectivity at multiple scales of space and time affects the spatial distribution of ecological and evolutionary processes (Gilarranz et al. 2017, Rayfield et al. 2023). Thus, there is a pressing need to facilitate the movements of multiple species at multiple spatial scales (Gonzalez et al., 2011; Rayfield et al., 2016; Thompson & Gonzalez 2017, Wood et al. 2022). The RE-Connect approach could be applied at several spatial scales to identify both short- and long-range connectivity priorities for distinct species (Albert et al., 2017; Fletcher et al., 2023).

### Conclusions

Despite its importance, assessments of ecological connectivity are often not considered in conservation management (Baguette & Van Dyck, 2007; Saura et al., 2018; Ward et al., 2020). The KM-GBF of the UN CBD lacks a headline indicator for ecological connectivity. In our study, we identified a key-set of connectivity indicators suitable for monitoring change to assess progress toward national and international targets (Gonzalez, Chase, & O’Connor, 2023; Jetz et al., 2019; Tittensor et al., 2014). Using our multi-indicator, multispecies approach and R-tool, we evaluated the status and decadal change of habitat connectivity for a set forest species with different habitat needs and dispersal capacities. We found that despite a trend of increasing forest habitat area from 2011 to 2021, connectivity indicators, as well as average forest patch sizes decreased, revealing the effects of ongoing habitat fragmentation across the St-Lawrence Lowlands in Quebec. Human land-use change and associated habitat fragmentation are among the most important drivers of biodiversity loss (Crooks et al., 2017; Haddad et al., 2015; Sala et al., 2000). At this time roughly half of the globally protected areas are also connected (Saura et al., 2018; Ward et al., 2020; 9.7% out of 16.6 % of the land on Earth). Efforts to meet the targets of the KM-GBF have highlighted the need for connectivity conservation from local to global scales. This will require a rapid prioritization of areas for conservation actions and assessment of the efficacy of alternative conservation scenarios for the coming decades. The research presented here and the RE-Connect approach and R-tool can help to support conservation management with quantitative indicators to monitor connectivity change at different scales for a wide range of species.

## Acknowledgements

We acknowledge the funding and the financial support by McGill University, Habitat (habitat-nature.com) and the Mitacs foundation (IT19834). We also thank Environment Climate Change Canada (ECCC) and Dr. Amanda Martin for their financial support (GCXE22S043) and stimulating discussions.

## Supporting Information

Additional supporting information may be found in the online version of the article at the publisher’s website.

