## Supporting Information for "Rapid evaluation of habitat connectivity change to safeguard multispecies persistence in human-transformed landscapes"

Appendix S1

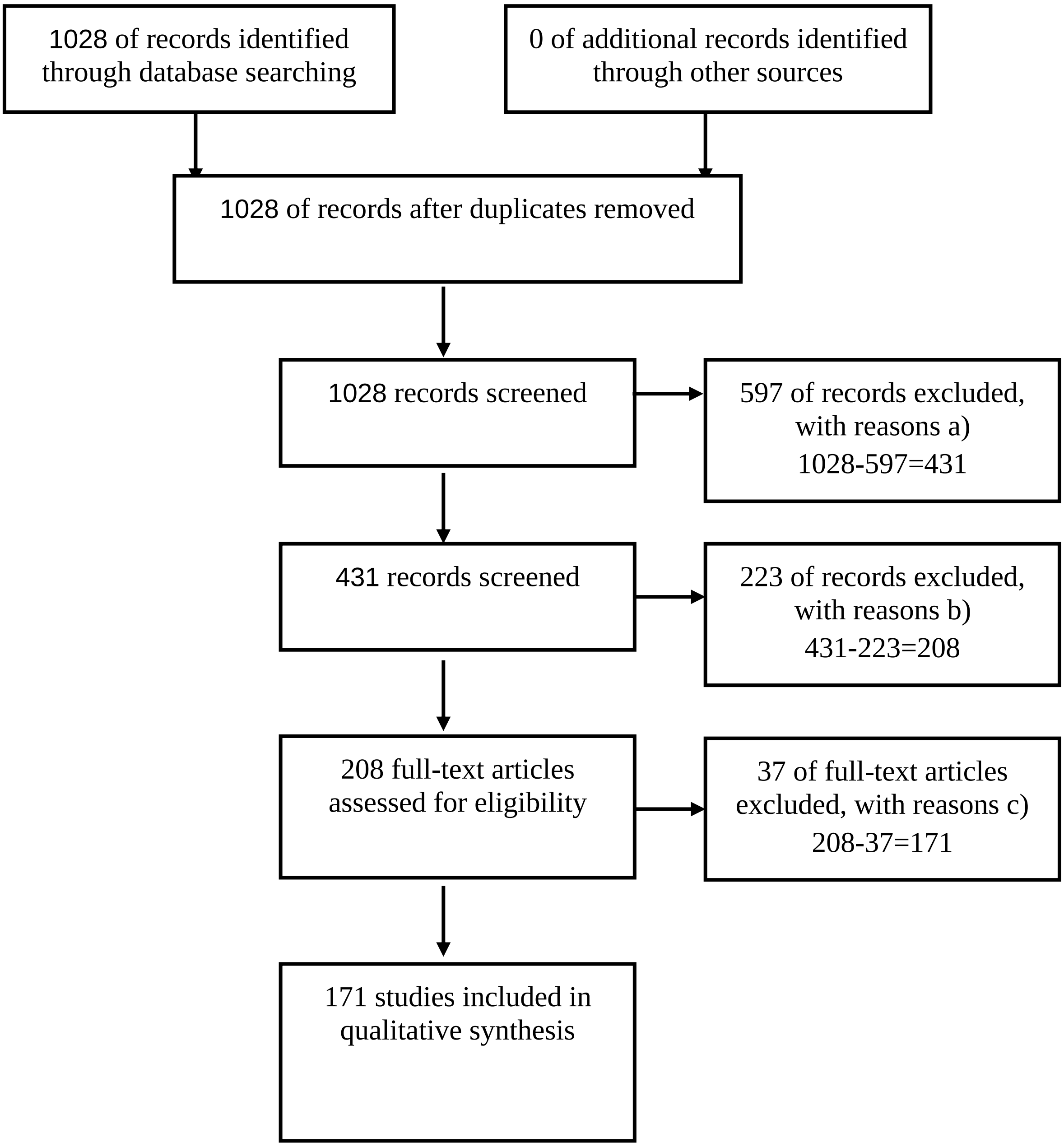

**Supporting Figure 1. PRISMA flow diagram (Liberati et al., 2009) describing the selection procedure in our literature review.** Using our search string (cf. Methods), we searched the Scopus and Web of Science databases on January 24th, 2022, and found 1028 journal articles published since January 1, 2000. Reasons to exclude studies: **a)** articles that did not mention either connectivity or monitoring in their title, **b)** articles that i) were not concerned with habitat connectivity, ii) were not primary research articles, iii) did not use any indicator of connectivity, and iv) were focused on a single species only, whereby we kept articles that used a single species as umbrella species. **c)** articles that did not align with our criteria in a) or b).

### Appendix S2

**Supporting Table 1. Criteria for the selection of key-connectivity indicators[1].**

| Nr | Abbreviation | feasibility (commonness, ease of computation) | relevance (alignment with target) | scalability | sensitivity | scale, interpretation |
|---|---|---|---|---|---|---|
| **1** | **MPC** | high, >3 case studies, binary habitat distribution maps | species persistence | yes | yes | landscape level species persistence |
| **2** | **ECA** | high, >3 case studies, binary habitat distribution maps | area-based conservation | yes | yes | landscape level amount of connected area |
| **3** | **$ECA_{Ap}$** | low, 0 case studies, binary habitat distribution maps | area-based conservation | yes | yes | landscape level fraction of habitat that is connected |
| **4** | **$ECA_{AI}$** | high, >3 case studies, binary habitat distribution maps | area-based conservation | yes | yes | landscape level amount of habitat that is connected |
| **5** | **BC** | high, >3 case studies, binary habitat distribution maps | spatial prioritization | yes | yes | patch level importance for short and long-range movement |
| **6** | **ND** | high, >3 case studies, binary habitat distribution maps | spatial prioritization | yes | yes | patch level importance for short-range movement |
| **7** | **invCR** | medium, 1 case study, resistance map | spatial prioritization | yes | yes | pixel level contribution to short- and long-range movement |
| **8** | **$I_v$** | high, >3 case studies, binary habitat distribution maps | spatial prioritization | yes | yes | patch level contribution to landscape level connectivity index I |
| **9** | **$MPC_i$** | high, >3 case studies, binary habitat distribution maps | spatial prioritization | yes | yes | patch level contribution to landscape level metapopulation capacity |

[1]We selected key-connectivity indicators identified in our literature review based on feasibility, relevance, scalability and sensitivity criteria established in the Essential Biodiversity Variable's framework (EBV) (Balvanera et al., 2022; Jetz et al., 2019; Pereira et al., 2013). The list of selected indicators includes graph-based metrics (Minor & Urban, 2008) such as the effectively connected habitat area (Saura, Bastin, Battistella, Mandrici, & Dubois, 2017) (ECA), and the importance of stepping stones (Albert et al. 2017) (betweenness centrality [BC] and node degree [ND]), resistance-distance derived proxies of landscape traversability (Albert et al., 2017; Chubaty, Galpern, & Doctolero, 2020; Shahnaseri et al., 2019) (invCR), and metapopulation-model derived long-term persistence of species (Hanski & Ovaskainen, 2000; Schnell et al., 2013) (MPC). These connectivity indicators also span different and complementary scales of spatial organization (Fletcher et al., 2023). For example, ECA and MPC are typically assessed at the scale of a landscape, BC and ND are typically assessed at the scale of habitat patches, and finally, resistance-based proxies of landscape traversability such as invCR are assessed at the pixel (or plot) level (Fletcher et al., 2023). We assessed the sensitivity of selected indicators to changes in habitat area and fragmentation by means of simulated and real-world landscapes (see Figure 5 and Appendix S3).

#### Appendix S3

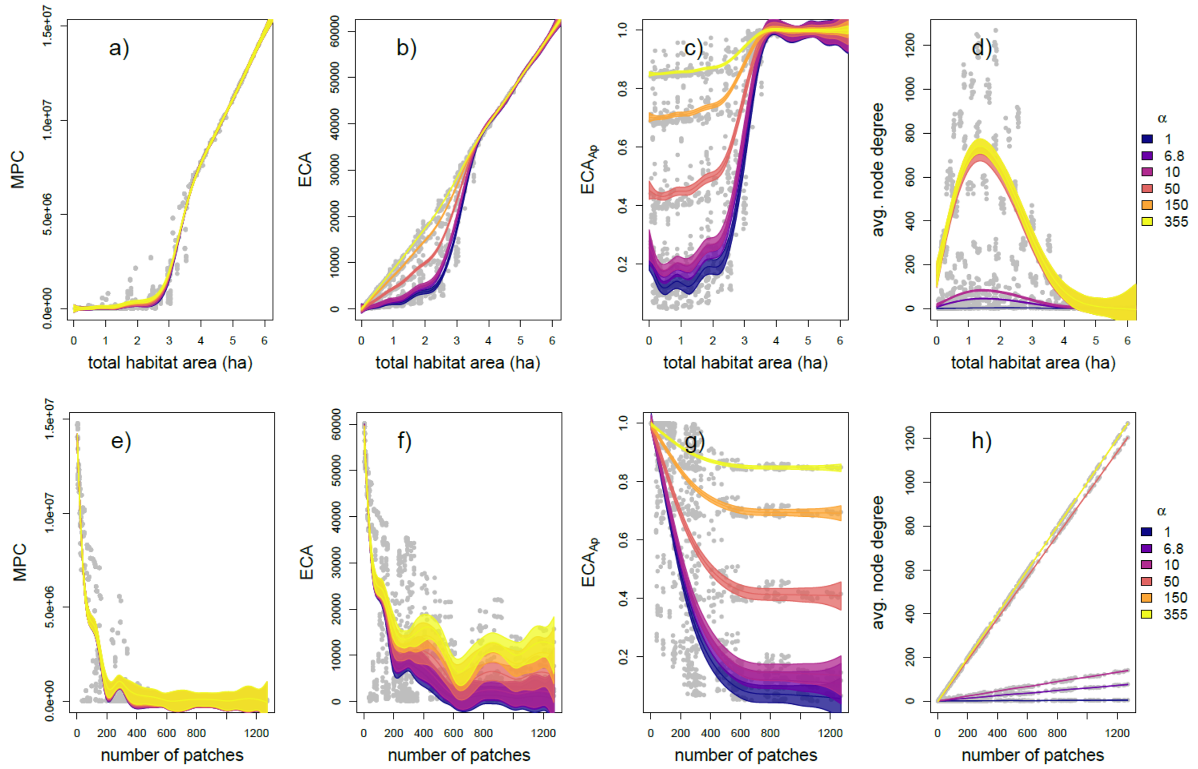

**Supporting Figure 2. Change of multiple connectivity dimensions with total habitat area (a-d) and habitat fragmentation (approximated by the number of habitat patches, e-h) in simulated landscapes (Saura & Martínez-Millán, 2000).** Landscape-level connectivity values (Table 1) were calculated using the RE-Connect R-tool and a library of 270 simulated landscapes varying in habitat area and fragmentation for different dispersal capacities (Methods). Compared with landscape size (250 x 250 cells), a gap crossing distance of 6.8 corresponds most closely to the maximum of the gap crossing distance among our species (236 m) in the 75 $km^2$ moving windows (~1/36 of the landscape side length, cf. Methods). Relationships among variables were estimated and predicted (fitted values ± standard error) using the smooth spline function in the npreg R-package (Helwig, 2021). **a & e)** MPC: metapopulation capacity, **b & f)** ECA: equivalent connected area index, **c & g)** $ECA_{Ap}$: fraction of habitat that is connected, **d & h)** average node degree of habitat patches in the landscape.

Appendix S4

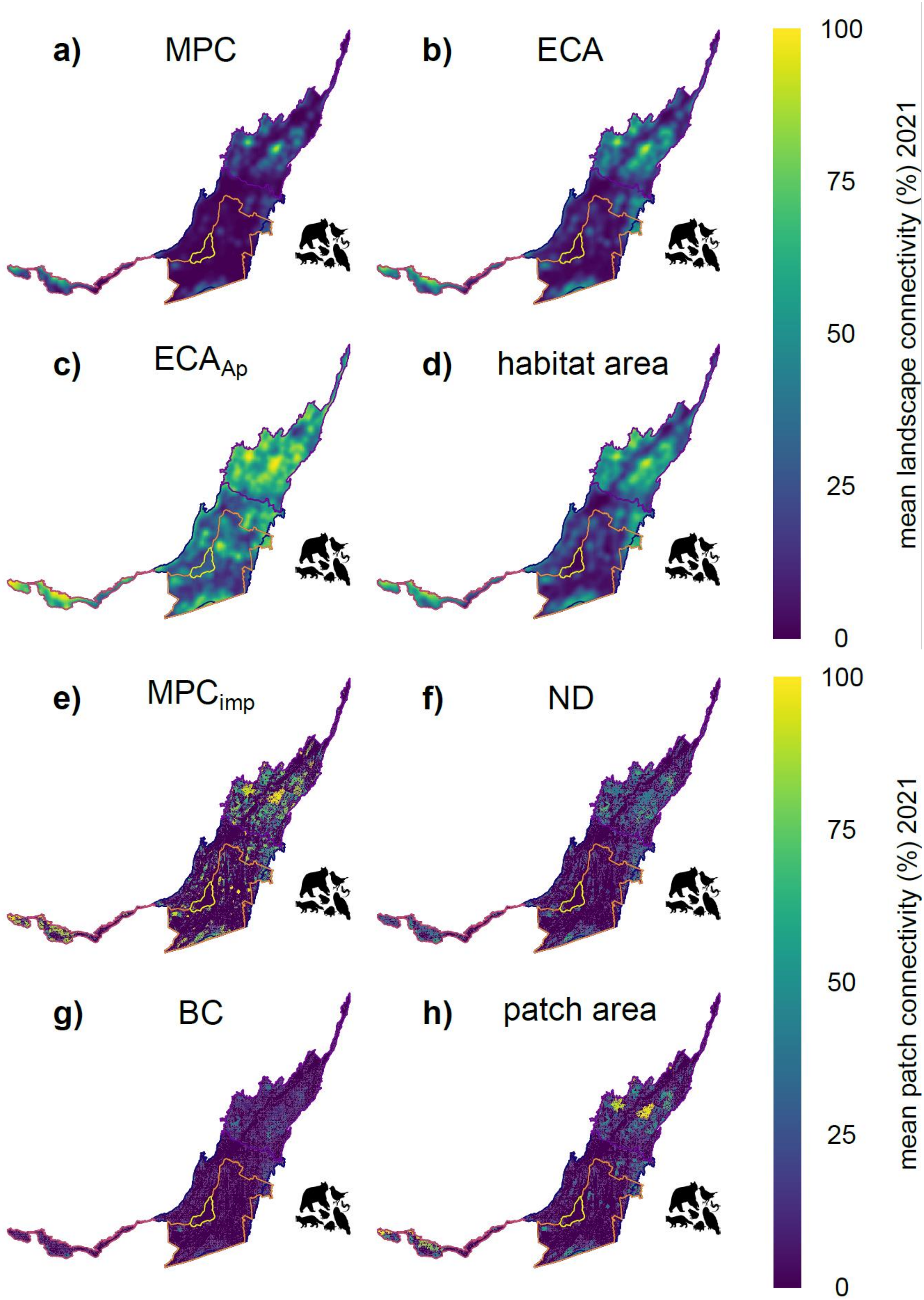

**Supporting Figure 3. State of multispecies connectivity at landscape-level (a-d) and patch-level (e-h) across the St-Lawrence Lowlands in 2021.** Using RE-Connect, we mapped the spatial distribution of multiple connectivity dimensions (Table 1), normalized and averaged across seven ecoprofile species representing forest connectivity needs (cf. Methods and Table 2). **a)** MPC: metapopulation capacity, **b)** ECA: equivalent connected area index, based on the probability of connectivity index (PC), **c)** $ECA_{Ap}$: fraction of habitat that is connected (ECA divided by the total amount of habitat area), **d)** habitat area: area of habitat, **e)** $MPC_{imp}$: metapopulation capacity patch importance, i.e. the contribution of a habitat patch to landscape-level metapopulation capacity, **f)** ND: node degree of a focal patch, i.e. the number of other habitat patches connected to the focal patch, **g)** BC: betweenness centrality of a focal patch, i.e. the number of shortest paths between all other pairs of habitat patches in the landscape that go through the focal patch, **h)** patch area: area of the focal patch.

Appendix S5

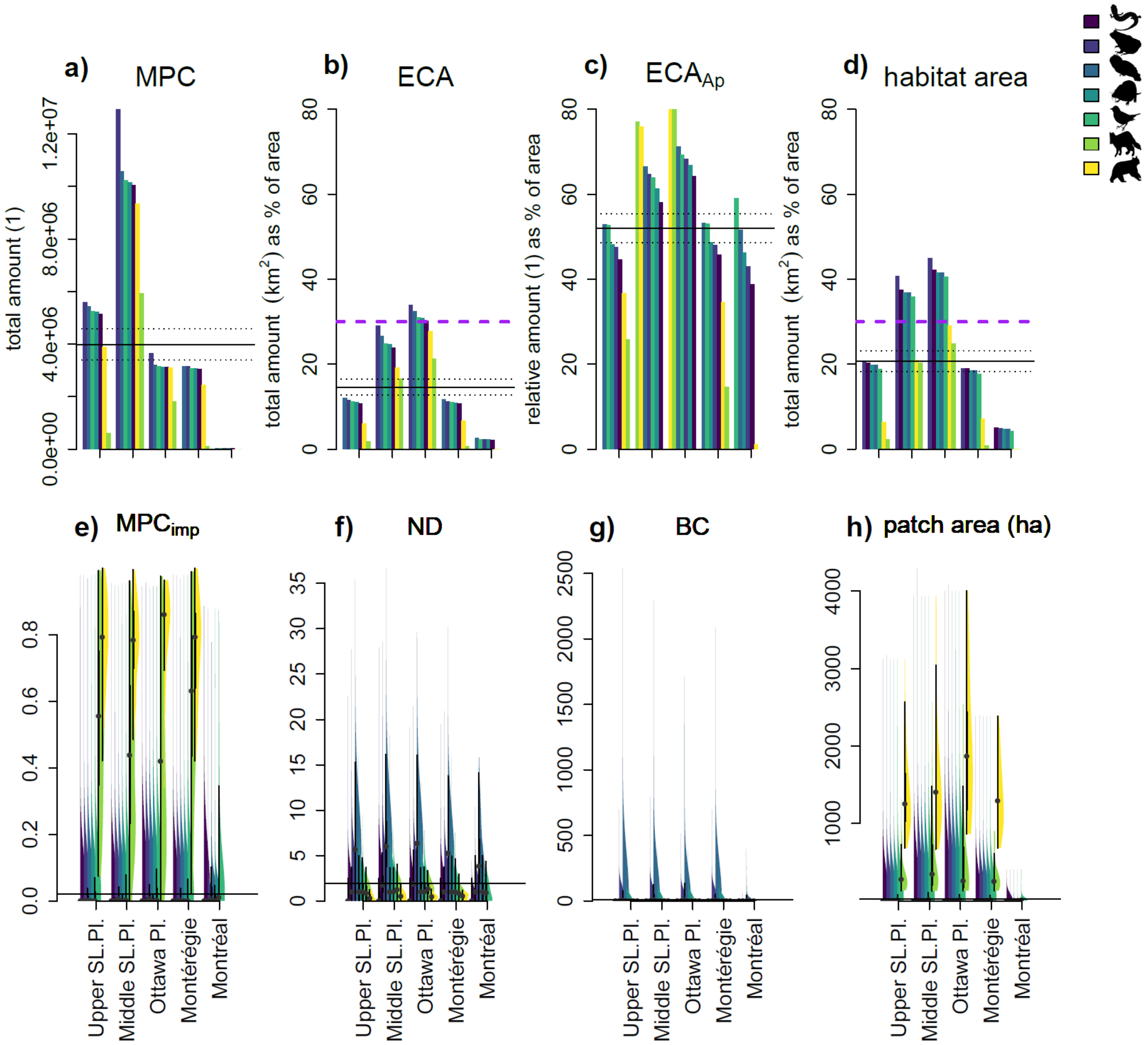

**Supporting Figure 4. Status (2021) of multispecies connectivity at landscape-level (a-d) and patch-level (e-h) for 7 species in 5 different regions in the St-Lawrence Lowlands.** **a)** MPC: metapopulation capacity, **b)** ECA: equivalent connected area index, based on the probability of connectivity index (PC), **c)** $ECA_{Ap}$: fraction of habitat that is connected (ECA divided by the total amount of habitat area), **d)** habitat area: species-specific area of habitat in the landscape, **e)** $MPC_{imp}$: metapopulation capacity patch importance, i.e. the contribution of a habitat patch to landscape-level metapopulation capacity, **f)** ND: node degree of a focal patch: the number of other habitat patches connected to the focal patch, **g)** BC: betweenness centrality of a focal patch: the number of shortest paths between all other pairs of habitat patches in the landscape that go through the focal patch, **h)** patch area (ha): habitat patch area in hectares. We multiplied RE-Connect-derived landscape-connectivity values with the area they cover in $km^2$ and consequently summed the results for each region. We scaled connectivity values by the area of the RE-Connect moving window size ($8{,}700^2 = 75.69\ km^2$) prior to the summation if necessary, i.e. in the cases of MPC, ECA and habitat area. We divided the results by the total area of each region (in $km^2$) and multiplied by 100, in order to get relative area estimates in % (y-axis, b-d). For the patch-level connectivity indicators, we extracted the median and interquartile range (violin plots, values per patch on y-axes e-h). Black horizontal bars indicate the mean and s.e. of connectivity values across all regions and species. Purple horizontal bars indicate a 30% (connected-) habitat area target as described in Target 3 of the Post-2020 Global Biodiversity Framework of the Convention on Biological Diversity (CBD, 2021). See Appendix S6 and S7 for more details.

#### Appendix S6

**Supporting Table 2. Total amount (2021) and change in amount (Δ, 2011-2021) of landscape-level forest connectivity for 7 ecoprofile species and 5 regions in the St-Lawrence Lowlands[1].**

| **region** | **species** | **MPC** total amount (1) 2021 | **ΔMPC** total amount (1) 2011-2021 | **ECA** total amount ($km^2$) 2021 | **ΔECA** total amount ($km^2$) 2011-2021 | **ECAf** fraction of area (%) 2021 | **ΔECAf** fraction of area (%) 2011-2021 | **$ECA_{Ap}$** fraction of area (%) 2021 | **$ΔECA_{Ap}$** fraction of area (%) 2011-2021 | **habitat area** total amount ($km^2$) 2021 | **Δhabitat area** total amount ($km^2$) 2011-2021 | **habitat area_f** fraction of area (%) 2021 | **Δhabitat area_f** fraction of area (%) 2011-2021 |
|---|---|---|---|---|---|---|---|---|---|---|---|---|---|
| Upper SL.Pl. | BLBR | 5218326 | -1452345 | 1923 | -113 | 11.11 | -0.65 | 48.16 | -6.05 | 3435 | 133 | 19.84 | 0.77 |
| Upper SL.Pl. | MAAM | 615606 | -718270 | 302 | -214 | 1.75 | -1.23 | 25.88 | -10.29 | 388 | -249 | 2.24 | -1.44 |
| Upper SL.Pl. | PLCI | 5149670 | -1483233 | 1859 | -131 | 10.74 | -0.76 | 44.56 | -8.07 | 3522 | 209 | 20.34 | 1.21 |
| Upper SL.Pl. | RASY | 5598822 | -1270356 | 2008 | -53 | 11.60 | -0.31 | 47.59 | -6.55 | 3562 | 232 | 20.57 | 1.34 |
| Upper SL.Pl. | SEAU | 5252811 | -1442902 | 1939 | -115 | 11.20 | -0.66 | 52.66 | -3.19 | 3275 | 17 | 18.92 | 0.10 |
| Upper SL.Pl. | STVA | 5441259 | -1430513 | 2095 | -94 | 12.10 | -0.54 | 52.93 | -5.54 | 3435 | 133 | 19.84 | 0.77 |
| Upper SL.Pl. | URAM | 3885946 | -1693446 | 1042 | -256 | 6.02 | -1.48 | 36.72 | -7.54 | 1113 | -230 | 6.43 | -1.33 |
| Middle SL.Pl. | BLBR | 10159294 | -3678927 | 2749 | -236 | 24.64 | -2.11 | 61.24 | -5.15 | 4110 | 33 | 36.85 | 0.29 |

| | | | | | | | | | | | | | |
|---|---|---|---|---|---|---|---|---|---|---|---|---|---|
| Middle SL.Pl. | MAAM | 5934772 | 877180 | 1840 | 252 | 16.50 | 2.26 | 76.95 | 3.34 | 2258 | 214 | 20.24 | 1.92 |
| Middle SL.Pl. | PLCI | 10042053 | -3719929 | 2664 | -253 | 23.88 | -2.27 | 58.01 | -6.63 | 4181 | 97 | 37.48 | 0.87 |
| Middle SL.Pl. | RASY | 12939631 | -1990065 | 3241 | 115 | 29.05 | 1.03 | 64.62 | -2.44 | 4549 | 335 | 40.78 | 3.00 |
| Middle SL.Pl. | SEAU | 10221514 | -3665987 | 2773 | -240 | 24.86 | -2.15 | 63.93 | -3.64 | 4006 | -50 | 35.91 | -0.44 |
| Middle SL.Pl. | STVA | 10571627 | -3623517 | 2968 | -212 | 26.61 | -1.90 | 66.57 | -4.69 | 4110 | 33 | 36.85 | 0.29 |
| Middle SL.Pl. | URAM | 9335380 | -4052682 | 2135 | -462 | 19.14 | -4.14 | 75.87 | -2.99 | 2341 | -413 | 20.98 | -3.70 |
| Ottawa Pl. | BLBR | 3137986 | -557338 | 685 | -47 | 30.82 | -2.12 | 66.85 | -3.92 | 924 | -26 | 41.55 | -1.18 |
| Ottawa Pl. | MAAM | 1807935 | 154435 | 473 | 22 | 21.28 | 1.01 | 80.35 | 3.29 | 551 | -9 | 24.77 | -0.42 |
| Ottawa Pl. | PLCI | 3120276 | -560941 | 671 | -49 | 30.18 | -2.19 | 64.14 | -5.06 | 937 | -14 | 42.17 | -0.64 |
| Ottawa Pl. | RASY | 3639774 | -216113 | 753 | 1 | 33.89 | 0.04 | 68.28 | -2.78 | 999 | 26 | 44.95 | 1.18 |
| Ottawa Pl. | SEAU | 3147065 | -556533 | 688 | -49 | 30.96 | -2.19 | 69.27 | -2.62 | 901 | -43 | 40.53 | -1.92 |
| Ottawa Pl. | STVA | 3203250 | -559328 | 723 | -48 | 32.51 | -2.17 | 71.19 | -3.82 | 924 | -26 | 41.55 | -1.18 |

| | | | | | | | | | | | | | |
|---|---|---|---|---|---|---|---|---|---|---|---|---|---|
| Ottawa Pl. | URAM | 3090734 | -574885 | 617 | -62 | 27.74 | -2.80 | 86.06 | -3.97 | 645 | -63 | 29.04 | -2.83 |
| Montérégie | BLBR | 3061669 | -451353 | 1043 | -17 | 10.96 | -0.18 | 48.72 | -5.88 | 1763 | 103 | 18.53 | 1.09 |
| Montérégie | MAAM | 122200 | -153307 | 68 | -59 | 0.71 | -0.62 | 14.63 | -10.30 | 85 | -66 | 0.89 | -0.69 |
| Montérégie | PLCI | 3035636 | -457852 | 1016 | -23 | 10.67 | -0.24 | 45.72 | -7.48 | 1805 | 141 | 18.97 | 1.49 |
| Montérégie | RASY | 3144562 | -447489 | 1063 | -4 | 11.17 | -0.04 | 48.00 | -6.58 | 1807 | 139 | 18.99 | 1.46 |
| Montérégie | SEAU | 3074577 | -449320 | 1049 | -19 | 11.03 | -0.20 | 53.05 | -3.20 | 1681 | 41 | 17.66 | 0.44 |
| Montérégie | STVA | 3148773 | -444395 | 1121 | -7 | 11.78 | -0.07 | 53.15 | -5.47 | 1763 | 103 | 18.53 | 1.09 |
| Montérégie | URAM | 2446303 | -445314 | 631 | -25 | 6.63 | -0.27 | 34.61 | -4.55 | 679 | 2 | 7.14 | 0.02 |
| Montréal | BLBR | 20574 | 2414 | 15 | 2 | 2.36 | 0.35 | 46.30 | -8.94 | 30 | 4 | 4.72 | 0.69 |
| Montréal | MAAM | 0.00 | -220 | 0.00 | -0.17 | 0.00 | -0.03 | 0.00 | -1.52 | 0.00 | -0.18 | 0.00 | -0.03 |
| Montréal | PLCI | 19429 | 1832 | 14 | 2 | 2.20 | 0.29 | 38.82 | -11.23 | 32 | 6 | 5.07 | 0.94 |
| Montréal | RASY | 20655 | 2456 | 15 | 2 | 2.38 | 0.36 | 42.92 | -9.66 | 31 | 5 | 4.94 | 0.85 |

| | | | | | | | | | | | | | |
|---|---|---|---|---|---|---|---|---|---|---|---|---|---|
| Montréal | SEAU | 21055 | 2656 | 15 | 2 | 2.38 | 0.36 | 58.97 | 0.68 | 27 | 3 | 4.23 | 0.41 |
| Montréal | STVA | 22923 | 3369 | 17 | 3 | 2.68 | 0.44 | 51.56 | -7.61 | 30 | 4 | 4.72 | 0.69 |
| Montréal | URAM | 147 | -7887 | 0.09 | -2.60 | 0.01 | -0.41 | 1.15 | -5.63 | 0.09 | -2.60 | 0.01 | -0.41 |

[1]These results correspond to data shown in Figure 4a-d and Appendix S5a-d. For recent changes in landscape-level connectivity (**Δ**, 2011-2021), decreases are highlighted in red and increases highlighted in blue. MPC: metapopulation capacity, ECA: equivalent connected area index, based on the probability of connectivity index (PC), ECAf: ECA as % of the area in a region, $ECA_{Ap}$: fraction of habitat that is connected (ECA divided by the total amount of habitat area), as % of the area in a region, habitat area: species-specific area of habitat in the region, habitat area_f: species-specific habitat area as % of the area in a region. PLCI: Red-back salamander, RASY: Wood frog, STVA: Barred Owl, BLBR: Northern short-tailed shrew, SEAU: Ovenbird, MAAM: American marten, URAM: Black bear. Computation of total amounts of connectivity was done in each region by a simple multiplication of RE-Connect-derived landscape-level connectivity values with the area they cover in $km^2$ and consequent summation. Note that we scaled connectivity values by the area of the RE-Connect moving window size ($8{,}700^2$ = 75.69 $km^2$) prior to the summation if necessary, i.e. in the cases of MPC, ECA and habitat area. Species that meet a 30% area-based conservation target for ECA and habitat area are highlighted in yellow. See Table 1 for more details on connectivity indicators.

Appendix S7

**Supporting Table 3. State (2021) and change (2011-2021) of patch-level forest connectivity for 7 ecoprofile species and 5 regions in the St-Lawrence Lowlands[1].**

| region | species | nr. patches 2021 | BC mn±sd 2021 | ND mn±sd 2021 | MPCimp mn±sd 2021 | ECAimp mn±sd 2021 | Patch area (ha) mn±sd 2021 | Δ nr. patches 2011-2021 | Δ BC mn±se 2011-2021 | Δ ND mn±se 2011-2021 | Δ MPCimp mn±se 2011-2021 | Δ ECAimp mn±se 2011-2021 | Δ Patch area (ha) mn±se 2011-2021 |
|---|---|---|---|---|---|---|---|---|---|---|---|---|---|
| Upper SL.Pl. | BLBR | 11759 | 1.7±9.71 | 1.4±1.44 | 0.02±0.1 | 1.29±5.9 | 26.69±126.38 | 6627 | 1.11±0.1 | 0.39±0.02 | -0.02±0 | -1.32±0.13 | -24.98±2.9 |
| Upper SL.Pl. | MAAM | 101 | 0.61±1.4 | 1.02±0.88 | 0.55±0.26 | 40.11±25.2 | 350.61±240.97 | -33 | 0.04±0.17 | 0.09±0.12 | 0±0.03 | -1.1±3.44 | -46.48±37.54 |
| Upper SL.Pl. | PLCI | 30908 | 0±0 | 0.59±0.98 | 0.01±0.06 | 0.47±3.66 | 10.48±79.13 | 23575 | NA | 0.21±0.01 | -0.02±0 | -1.29±0.09 | -25.95±1.93 |
| Upper SL.Pl. | RASY | 18568 | 2.9±18.66 | 1.72±1.78 | 0.02±0.08 | 0.83±4.8 | 17.72±106.61 | 12469 | 2.08±0.15 | 0.61±0.02 | -0.02±0 | -1.37±0.11 | -26.23±2.42 |
| Upper SL.Pl. | SEAU | 4451 | 1.17±3.9 | 1.25±1.15 | 0.06±0.16 | 3.38±9.3 | 66.51±198.58 | 1034 | 0.61±0.07 | 0.19±0.02 | -0.01±0 | -0.55±0.23 | -9.55±4.91 |
| Upper SL.Pl. | STVA | 11759 | 39.24±112.44 | 6.36±4.06 | 0.03±0.1 | 1.43±5.91 | 26.69±126.38 | 6627 | 26.43±1.14 | 2.44±0.05 | -0.03±0 | -1.46±0.13 | -24.98±2.9 |
| Upper SL.Pl. | URAM | 65 | 0.07±0.13 | 0.42±0.4 | 0.78±0.14 | 68.07±23.6 | 1383.59±485.86 | 23 | NA | 0.2±0.07 | -0.1±0.02 | -22.22±3.85 | -320.85±129.46 |
| Middle SL.Pl. | BLBR | 7962 | 2.49±11.46 | 1.63±1.59 | 0.02±0.1 | 1.22±6.32 | 39.07±198.79 | 5008 | 1.91±0.13 | 0.54±0.03 | -0.02±0 | -1.19±0.18 | -35.01±5.43 |

| | | | | | | | | | | | | | |
|---|---|---|---|---|---|---|---|---|---|---|---|---|---|
| Middle SL.Pl. | MAAM | 289 | 0.89±1.58 | 1.36±0.9 | 0.44±0.25 | 28.39±21.01 | 540.89±484.5 | -18 | -0.16±0.14 | -0.09±0.07 | 0.03±0.02 | 2.5±1.69 | 83.69±38.12 |
| Middle SL.Pl. | PLCI | 24512 | 0±0 | 0.73±1.19 | 0.01±0.05 | 0.38±3.62 | 13.1±115.02 | 20164 | NA | 0.34±0.01 | -0.02±0 | -1.2±0.12 | -37.97±3.5 |
| Middle SL.Pl. | RASY | 11246 | 3.77±18.63 | 1.88±1.9 | 0.02±0.08 | 0.86±5.48 | 29.35±186.68 | 7758 | 3.02±0.18 | 0.73±0.03 | -0.02±0 | -1.16±0.15 | -33.35±4.6 |
| Middle SL.Pl. | SEAU | 2821 | 1.97±5.61 | 1.54±1.23 | 0.06±0.16 | 3.41±10.24 | 104.97±322.28 | 726 | 1.16±0.12 | 0.25±0.03 | 0±0 | 0±0.3 | 2.77±9.13 |
| Middle SL.Pl. | STVA | 7962 | 55.97±130.03 | 6.76±4.08 | 0.03±0.1 | 1.33±6.38 | 39.07±198.79 | 5008 | 42.48±1.54 | 2.93±0.06 | -0.02±0 | -1.37±0.18 | -35.01±5.43 |
| Middle SL.Pl. | URAM | 86 | 0.09±0.18 | 0.58±0.48 | 0.78±0.12 | 66.33±21.8 | 1606.52±783.62 | 38 | 0.01±0.04 | 0.02±0.09 | -0.04±0.02 | -9.69±3.8 | -197.91±145.66 |
| Ottawa Pl. | BLBR | 1740 | 3.08±14.41 | 1.66±1.61 | 0.02±0.09 | 1.07±6.79 | 38.38±237.08 | 949 | 2.24±0.37 | 0.5±0.06 | -0.01±0 | -0.75±0.35 | -27.67±12.12 |
| Ottawa Pl. | MAAM | 76 | 0.78±1.27 | 1.31±0.83 | 0.43±0.29 | 28.59±23.41 | 557.27±615.26 | -8 | -0.51±0.26 | -0.21±0.14 | 0.09±0.05 | 6.55±3.52 | 97.23±87.54 |
| Ottawa Pl. | PLCI | 5028 | 0±0 | 0.68±1.14 | 0.01±0.06 | 0.36±4.01 | 13.59±140.66 | 3849 | NA | 0.24±0.03 | -0.01±0 | -0.82±0.22 | -30.94±7.52 |
| Ottawa Pl. | RASY | 2596 | 4.44±21.22 | 1.97±1.88 | 0.01±0.08 | 0.67±5.44 | 25.79±200.63 | 1645 | 3.5±0.44 | 0.71±0.05 | -0.01±0 | -0.78±0.27 | -28.48±9.28 |
| Ottawa Pl. | SEAU | 647 | 2.33±7.35 | 1.55±1.29 | 0.05±0.15 | 2.88±10.93 | 99.38±381.2 | 124 | 1.17±0.34 | 0.24±0.07 | 0±0.01 | 0.13±0.63 | 1.23±21.93 |
| Ottawa Pl. | STVA | 1740 | 56.68±128.16 | 6.96±4 | 0.03±0.09 | 1.16±6.82 | 38.38±237.08 | 949 | 36.73±3.45 | 2.42±0.13 | -0.02±0 | -0.84±0.35 | -27.67±12.12 |

| | | | | | | | | | | | | | |
|---|---|---|---|---|---|---|---|---|---|---|---|---|---|
| Ottawa Pl. | URAM | 20 | 0.06±0.15 | 0.5±0.47 | 0.81±0.15 | 77.76±19.47 | 1989.92±924.52 | 8 | 0.01±0.05 | -0.05±0.15 | -0.04±0.05 | -2.93±7.04 | -275.11±334.5 |
| Montérégie | BLBR | 6021 | 1.25±8.42 | 1.27±1.32 | 0.02±0.1 | 1.32±6.03 | 24.34±121.76 | 3443 | 0.81±0.12 | 0.36±0.03 | -0.03±0 | -1.56±0.2 | -25.04±3.88 |
| Montérégie | MAAM | 28 | 0.15±0.43 | 0.8±0.68 | 0.6±0.26 | 45.36±25.62 | 335.87±224.16 | -11 | -0.14±0.15 | 0.3±0.17 | -0.04±0.07 | -8.98±6.93 | -29.53±64.06 |
| Montérégie | PLCI | 15092 | 0±0 | 0.54±0.92 | 0.01±0.07 | 0.5±3.84 | 9.99±77.78 | 11755 | NA | 0.21±0.01 | -0.03±0 | -1.64±0.14 | -28.27±2.83 |
| Montérégie | RASY | 9362 | 2.08±14.57 | 1.55±1.59 | 0.02±0.08 | 0.85±4.89 | 16.12±100.93 | 6443 | 1.42±0.17 | 0.57±0.03 | -0.03±0 | -1.67±0.17 | -27.38±3.35 |
| Montérégie | SEAU | 2312 | 0.88±3.07 | 1.13±1.08 | 0.06±0.16 | 3.44±9.43 | 59.79±191.24 | 530 | 0.49±0.07 | 0.17±0.03 | -0.01±0.01 | -0.76±0.32 | -10.48±6.43 |
| Montérégie | STVA | 6021 | 29.08±95.33 | 5.77±3.6 | 0.03±0.1 | 1.46±6.02 | 24.34±121.76 | 3443 | 18.15±1.41 | 2.25±0.07 | -0.03±0 | -1.71±0.2 | -25.04±3.88 |
| Montérégie | URAM | 32 | 0.06±0.13 | 0.49±0.36 | 0.76±0.15 | 63.3±25.05 | 1390.53±472.04 | 10 | NA | 0.22±0.09 | -0.11±0.04 | -21.77±6.3 | -180.03±174.86 |
| Montréal | BLBR | 201 | 0.67±2.17 | 1.19±1.23 | 0.04±0.13 | 2.3±6.6 | 10.79±39.23 | 92 | 0.58±0.16 | 0.45±0.12 | -0.02±0.02 | -1.66±0.99 | -4.76±4.18 |
| Montréal | MAAM | 0 | NA | NA | NA | NA | NA | 0 | NA | NA | NA | NA | NA |
| Montréal | PLCI | 644 | 0±0 | 0.38±0.67 | 0.01±0.08 | 0.68±3.64 | 3.68±22.4 | 434 | NA | 0.17±0.04 | -0.02±0.01 | -1.39±0.5 | -4.63±1.92 |
| Montréal | RASY | 369 | 1.05±4.62 | 1.44±1.49 | 0.02±0.1 | 1.26±4.94 | 6.22±29.43 | 218 | 0.88±0.25 | 0.63±0.11 | -0.02±0.01 | -1.62±0.69 | -5.22±2.79 |

| | | | | | | | | | | | | | |
|---|---|---|---|---|---|---|---|---|---|---|---|---|---|
| Montréal | SEAU | 63 | 0.55±1.27 | 1±1.11 | 0.13±0.22 | 6.98±11.07 | 30.03±66.45 | 15 | NA | 0.31±0.17 | -0.01±0.04 | -1.2±2.31 | -2.65±10.5 |
| Montréal | STVA | 201 | 21.7±53.53 | 4.8±3.71 | 0.06±0.13 | 2.59±6.81 | 10.79±39.23 | 92 | 17.68±3.89 | 1.74±0.34 | -0.02±0.02 | -1.72±0.99 | -4.76±4.18 |
| Montréal | URAM | 0 | NA | NA | NA | NA | NA | 0 | NA | NA | NA | NA | NA |

[1]These results correspond to data shown in Figure 4e-h and Appendix S5e-h. For recent changes in connectivity (**Δ**, 2011-2021), decreases are highlighted in red and increases highlighted in blue. To quantify recent change of connectivity indicators we used two-sided Welch t-tests in each region of interest. BC: betweenness centrality of a focal patch: the number of shortest paths between all other pairs of habitat patches in the landscape that go through the focal patch, ND: node degree of a focal patch: the number of other habitat patches connected to the focal patch, $MPC_{imp}$: metapopulation capacity patch importance, i.e. the contribution of a habitat patch to landscape-level metapopulation capacity, $ECA_{imp}$: equivalent connected area patch importance, i.e. the contribution of a habitat patch to landscape-level ECA, Patch area (ha): habitat patch area in hectares. See Table 1 for more details on connectivity indicators. mn: mean, sd: standard deviation, se: standard error of the mean estimated difference in the Welch t-tests. PLCI: Red-back salamander, RASY: Wood frog, STVA: Barred Owl, BLBR: Northern short-tailed shrew, SEAU: Ovenbird, MAAM: American marten, URAM: Black bear.
